# The *Staphylococcus aureus* prophage-encoded SSBP attenuates virulence and enhances IL-6-mediated macrophage clearance

**DOI:** 10.1101/2025.03.13.643069

**Authors:** Nour Ahmad-Mansour, Cynthia N. Abi Najem, Marianne Martin, Lucile Plumet, Sylvaine Huc-Brandt, Madjid Morsli, Patrice François, Albert Sotto, Catherine Dunyach-Remy, Jean-Philippe Lavigne, Virginie Molle

## Abstract

Pathogens often manipulate host immune responses to promote infection. Here, we describe a novel mechanism by which a secreted prophage-encoded single-stranded DNA-binding protein (SSBP) attenuates *Staphylococcus aureus* virulence while enhancing host immune defenses. SSBP, encoded by the ROSA-like prophage in a colonizing, reduced-virulence strain of *S. aureus* (NSA1385), significantly increases bacterial susceptibility to macrophage-mediated killing *in vitro* and reduces pathogenicity in a zebrafish infection model. Treatment with purified recombinant SSBP decreases bacterial survival within macrophages and demonstrates therapeutic potential. Notably, SSBP activates macrophages via interleukin-6 (IL-6), a pro-inflammatory cytokine that promotes bacterial clearance and macrophage activation. These findings uncover an unexpected prophage-derived mechanism in a colonizing *S. aureus* strain that attenuates virulence while stimulating host immunity, offering promising avenues for anti-infective therapies and immunomodulatory strategies.

## INTRODUCTION

*Staphylococcus aureus* is a major opportunistic pathogen responsible for a wide range of infections, from benign skin lesions to life-threatening systemic conditions such as pneumonia, sepsis, and osteomyelitis (Tong *et al*., 2015; Cheung *et al*., 2021). Its ability to colonize both community and healthcare settings has made it a leading cause of morbidity and mortality worldwide. The increasing antibiotic resistance of this pathogen, particularly with the emergence of methicillin-resistant *S. aureus* (MRSA), has exacerbated the challenge of effective treatment, highlighting an urgent need for alternative therapeutic strategies (Corona and Guo, 2016; Vestergaard *et al*., 2019). *S. aureus* strains are especially problematic as they not only resist frontline antibiotics but also exhibit a remarkable capacity to evade host immune defenses, allowing them to establish persistent and recurrent infections. Thus, understanding the molecular mechanisms underlying *S. aureus* pathogenicity and its interactions with the host immune system is essential for developing innovative therapeutic strategies. Given that *S. aureus* colonizes approximately 30% of the human population, particularly in the nasal cavity, this commensal state plays a key role in shaping its relationship with the host. Studying the fine balance between colonization and infection is crucial to deciphering how *S. aureus* transitions from a harmless commensal to an opportunistic pathogen, evades immune responses, and establishes persistent infections.

A key factor contributing to the success of *S. aureus* as a pathogen is its exceptional genetic plasticity, which enables it to acquire new virulence and resistance traits through horizontal gene transfer. Mobile genetic elements, such as plasmids, transposons, staphylococcal pathogenicity islands, staphylococcal cassette chromosomes, and prophages (stable inserted bacteriophages or lysogenic phages), play central roles in this process (Malachowa and DeLeo, 2010; Hiramatsu *et al*., 2013; Lebeurre *et al*., 2019). Among these elements, prophages have been particularly well-studied for their ability to integrate into the bacterial genome and disseminate genes encoding virulence factors and toxins that enhance bacterial survival and immune evasion. Notable examples include diphtheria toxin in *Corynebacterium diphtheriae*, botulinum toxin in *Clostridium botulinum*, and Shiga toxin in *Escherichia coli* (Feiner *et al*., 2015). In *S. aureus*, prophage-encoded immune evasion clusters (IEC) and toxins, such as staphylococcal enterotoxins, further exacerbate its ability to circumvent host defenses, disrupt immune responses, and promote persistent infections (Tuffs *et al*., 2018).

Our findings, however, challenge the classical understanding of prophages as virulence enhancers. Our previous studies identified a *S. aureus* strain (NSA1385) isolated from a diabetic foot ulcer (DFU) that harbors a ROSA-like prophage associated with reduced virulence, an effect that is independent of the prophage insertion site (Messad *et al*., 2015; Rasigade *et al*., 2016; Ahmad-Mansour *et al*., 2023). DFUs represent a unique clinical niche where *S. aureus* can either persist as a colonizer or progress to invasive infection, such as osteomyelitis. Interestingly, *S. aureus* NSA1385 displayed markedly reduced pathogenicity in zebrafish infection models and impaired intracellular survival in osteoblasts compared to other virulent strains. These observations suggest that, under specific conditions, prophages can actively modulate *S. aureus* behavior to favor colonization over invasive disease. This challenges the classical view that prophages solely enhance virulence and raises important questions about the molecular mechanisms underlying these atypical effects. Interactions between *S. aureus* and host cells are central to its ability to colonize or invade tissues. While macrophages are critical components of host immunity, responsible for phagocytosis and pathogen clearance, *S. aureus* also demonstrates a remarkable ability to persist in non-phagocytic cells such as keratinocytes and osteoblasts (Löffler *et al*., 2014; Pidwill *et al*., 2021; Hommes and Surewaard, 2022). This intracellular persistence enables bacterial evasion of immune detection and antibiotic treatment, contributing to chronic and recurrent infections (Fraunholz and Sinha, 2012; Gunaratnam *et al*., 2019).

In this study, we sought to elucidate the molecular mechanisms by which the ROSA-like prophage attenuates *S. aureus* virulence and alters host-pathogen interactions. We report the identification of a prophage-encoded single-stranded DNA-binding protein (SSBP) that reduces *S. aureus* virulence by increasing bacterial susceptibility to macrophage-mediated killing through interleukin-6 (IL-6) activation. These findings reveal a previously unrecognized prophage-derived mechanism that modulates bacterial virulence and host immune responses, highlighting potential applications for both anti-infective strategies and therapeutic immune modulation.

## MATERIAL AND METHODS

### Bacterial Strains, Media, and Growth Conditions

The bacterial strains and plasmids used in this study are listed in Supplementary Table 1. *S. aureus* strains were cultivated in Tryptic Soy Broth (TSB) medium, and *E. coli* strains in Luria-Bertani (LB) medium, both at 37°C with shaking at 225 rpm. Ampicillin was used at a concentration of 100 µg/mL, and tetracycline at 10 µg/mL for strain construction and phenotypic selection. Bacterial growth was monitored using a microplate reader (Tecan, Model Spark, Grödig, Austria GmbH).

### Microarray Design

NSA1385 and Newman *S. aureus* strains were cultured overnight at 37°C in LB broth. Following dilution to an optical density of 600 nm (OD_600_) of 0.1, the bacterial cultures were grown to the mid-exponential phase. Cells were then harvested by centrifugation, and total RNA was extracted from the bacterial pellets using the RNeasy® Mini Kit (Qiagen, Courtaboeuf, France) following the manufacturer’s instructions. The extracted RNA was treated with RNase-free DNase (Qiagen) for 30 minutes at 37°C, followed by a second purification step. The purified RNA was stored at −80°C until further use. The microarray was manufactured through the *in situ* synthesis of 10,807 60-mer oligonucleotide probes (Agilent, Palo Alto, CA, USA), selected as previously described (Charbonnier *et al*., 2005). It covers 98% of all open reading frames (ORFs) annotated in various reference strains, including *S. aureus* N315, Mu50, MW2, COL, NCTC 8325, USA300, MRSA252, MSSA476, and Newman (Holden *et al*., 2004; Diep *et al*., 2006; Baba *et al*., 2008). Additionally, the microarray includes long oligonucleotide probes corresponding to ROSA-like prophage ORFs from NSA1385. For labeling, 5 µg of total RNA from NSA1385 was labeled with Cy3-dCTP, while 5 µg of total RNA from Newman was labeled with Cy5-dCTP, following the SuperScript II protocol (Thermo Fischer Scientific, France). The labeled RNAs were purified as previously described (Scherl *et al*., 2006). Hybridization of Cy5-labeled cDNA and Cy3-labeled cDNA was then performed, followed by scanning. Hybridization fluorescence intensities were quantified using Feature Extraction software (Agilent, version 8). Data from two independent biological experiments were analyzed using GeneSpring 8.0 (Silicon Genetics, Redwood City, CA, USA) with a 5% false discovery rate (FDR) (P-value cut-off: 0.05) and an arbitrary threshold of 1.5-fold to define significant differences in expression ratios.

### Construction of *S. aureus* Strain Overexpressing *ssbP*

The *ssbP* gene was amplified by PCR using primer pair #101/#102 (Sup. Table 2) and *S. aureus* NSA1385 chromosomal DNA as the template. The Spot fragment was amplified by PCR using primers #103/#104 and the pSpot2 vector as the template (Chromotek, Planegg, Germany). The *ssbP* plasmid was constructed using the NEB Gibson Assembly Kit (New England Biolabs, Ipswich, MA, USA). Purified *ssbP* PCR products were C-terminally fused to the Spot-tag and cloned into the EcoRI/BamHI-digested pTSS-Cm-Pcap vector (Schwendener and Perreten, 2015) via Gibson Assembly, generating the plasmid pTSS::ssb-P.

The plasmid was sequenced to confirm its validity, propagated in *E. coli* IM08B (Monk *et al*., 2015), and subsequently electroporated into the *S. aureus* Δ*rosa* strain.

### Cloning, Expression, and Purification of Recombinant SSBP

The *ssbP* gene was amplified by PCR using *S. aureus* NSA1385 chromosomal DNA as a template and primer pair #129/#130 (Supp. Table 2). The *ssbP* fragment was cloned into the NdeI/BamHI-digested pET19b vector using the NEB Gibson Assembly Kit (New England Biolabs), generating the plasmid pET19b::*ssbP*. For protein expression, *E. coli* BL21 Star cells harboring the pET19b::*ssbP* plasmid were cultured in LB medium at 37 °C until the culture reached an OD600 of 0.8. At this point, 0.5 mM isopropyl-β-D-thiogalactopyranoside (IPTG) was added to induce protein expression, and the culture was incubated for an additional 3 h at 37 °C. Protein purification was performed using TALON® metal affinity resin (Takara, Saint-Germain-en-Laye, France) following the manufacturer’s instructions. Briefly, bacterial cultures were centrifuged at 4,000 rpm for 15 minutes at 4 °C, and the resulting pellets were resuspended in lysis buffer (50 mM Tris-HCl [pH 7.4], 300 mM NaCl, 10% [vol/vol] glycerol, 10 mM imidazole), supplemented with DNase I and RNase A (final concentration 2 µg/ml each), and Complete™ protease inhibitor cocktail (Roche), to degrade contaminating nucleic acids and prevent protein degradation. Cells were lysed using a French pressure cell, and the lysates were centrifuged at 14,000 rpm for 30 min at 4 °C to remove cell debris. The supernatants were incubated with TALON® resin for one hour at 4 °C, then washed several times with wash buffer containing 50 mM Tris-HCl (pH 7.4), 300 mM NaCl, 10% (v/v) glycerol, and 50 mM imidazole. Elution was carried out with elution buffer containing 500 mM imidazole, 50 mM Tris-HCl (pH 7.4), 300 mM NaCl, and 10% (v/v) glycerol, to be stored at −20 °C.

### Electrophoretic Mobility Shift Assay (EMSA)

80-bp DYE682-labeled random primers (single-stranded DNA, ssDNA probe) (Eurofins Genomics, Nantes, France) were annealed by heating at 95°C for 5 minutes in 10 mM Tris-HCl (pH 8.0) and 20 mM NaCl, followed by cooling to 25°C to produce the double-stranded DNA (dsDNA) probe. The dsDNA and ssDNA probes were incubated with the indicated amounts of purified recombinant SSBP in 20 µL reactions containing EMSA buffer (20 mM HEPES [pH 8.0], 10 mM KCl, 2 mM MgCl₂, 0.1 mM EDTA, 0.1 mM dithiothreitol, 50 µg/mL bovine serum albumin) for 30 minutes. Samples were resolved on a 2.5% agarose gel run in cold TAE (Tris-Acetate-EDTA) buffer. Gel imaging was performed using the Odyssey Infrared Imaging System (LI-COR) on the 700 nm channel.

### Immunoprecipitation and Immunoblotting

*S. aureus* harboring the pTSS::*ssbP* plasmid was grown in TSB at 37°C with shaking (160 rpm) until reaching the exponential phase. Cultures were centrifuged for 10 minutes at 4,000 rpm, and Spot-fusion proteins were immunoprecipitated for 3 h at 4°C on a rotating wheel using anti-Spot beads (Chromotek). The bacterial pellet was washed twice with phosphate-buffered saline (PBS) and then resuspended in digestion buffer (50 mM Tris-HCl, 20 mM MgCl₂, 30% [wt/vol] raffinose, pH 7.5) supplemented with complete Mini EDTA-free protease inhibitors (Roche). Protein samples were resolved on an SDS-PAGE gel and transferred onto PVDF membranes for immunoblotting. The membrane was blocked overnight at 4°C in TBST (25 mM Tris-HCl, pH 7.6, 150 mM NaCl, 0.05% Tween-20) supplemented with 50% human serum (Sigma, ref. H3667) and 3% BSA. After blocking, the membrane was incubated for 1 h at room temperature with a monoclonal mouse anti-Spot antibody (1:5000, Chromotek) diluted in TBST containing 3% BSA. Following three washes with TBST, the membrane was incubated for 1 h at room temperature with an HRP-linked donkey anti-mouse secondary antibody (1:5000, Jackson ImmunoResearch, ref. 715-035-151) diluted in TBST with 3% BSA. After the membrane was washed three times with TBST, the HRP signal was revealed using the Pierce ECL Western Blotting Substrate and chemiluminescence was detected using Chemidoc imaging system (BioRad).

### Cell Culture and Infection Assays

The murine macrophage cell line RAW 264.7, derived from mouse leukemic monocyte macrophages (ATCC TIB-71), was cultured in Dulbecco’s Modified Eagle Medium (DMEM) supplemented with 10% fetal calf serum (FCS, Thermo Fisher Scientific) and maintained at 37°C with 5% CO₂. Similarly, the human leukemia monocytic cell line THP-1 was cultured in Roswell Park Memorial Institute (RPMI) medium supplemented with 10% FCS and maintained under the same conditions. THP-1 cells were differentiated into macrophages using phorbol myristate acetate (PMA) (50 ng/ml), as previously described (Tsuchiya *et al*., 1980; Takashiba *et al*., 1999). Macrophages were infected with *S. aureus* strains at a multiplicity of infection (MOI) of 20:1 (bacteria-to-cells ratio) for 1 h at 37°C with 5% CO₂. Prior to infection, *S. aureus* strains were cultured in TSB medium to the mid-exponential growth phase (OD_600_ = 0.6–0.9). After infection, cells were washed with PBS and treated with gentamicin (100 µg/mL) for 30 minutes to kill extracellular bacteria. Macrophages were then washed twice with PBS (T0) and incubated in fresh medium containing lysostaphin (5 µg/mL) for 5 and 24 h. Intracellular bacteria were quantified by lysing macrophages with 0.1% Triton X-100 in PBS, followed by serial dilution and plating on TSB agar for colony-forming unit (CFU) counting.

For macrophage infections treated with recombinant SSBP, RAW 264.7 cells were infected with *S. aureus* strains, and 3 µM recombinant SSBP was maintained in the culture medium throughout the experiment. A control group consisting only of elution buffer was included for comparison. Following the infection, an additional 3 µM of recombinant SSBP was freshly added to the culture medium along with lysostaphin, and the cells were incubated for 3 h.

HaCaT keratinocytes (Boukamp *et al*., 1988) and MC3T3-E1 osteoblasts (Sudo *et al*., 1983) were cultured in DMEM and α-MEM, respectively, supplemented with 10% FCS, 1% glutamine (Gibco™), and 0.5% penicillin-streptomycin antibiotics (Gibco™) in a humidified atmosphere at 37°C with 5% CO₂. *S. aureus* strains were cultivated in TSB medium to the mid-exponential growth phase, collected, resuspended in sterile PBS, and added to HaCaT and MC3T3-E1 cells at a MOI of 100:1. Extracellular bacteria were eliminated after 1.5 h of incubation by treating the cells with gentamicin. Infected cells were then washed twice with PBS and incubated in fresh medium containing lysostaphin for 5, 24, and 48 h. Intracellular bacteria were enumerated by lysing cells with 0.1% Triton X-100 in PBS, followed by serial dilution and plating on TSB agar plates. Lactate dehydrogenase (LDH) activity in cell culture supernatants was measured using the CyQUANT Cytotoxicity Assay Kit, following the manufacturer’s instructions.

### Zebrafish Infections

*S. aureus* strains were grown in TSB medium overnight at 37°C. Cultures were diluted 1:20 and cultivated in TSB medium until the mid-exponential growth phase (OD_600_ = 0.7-0.9) was attained. After centrifuging the bacteria for 10 minutes at 4,000 rpm, they were resuspended in PBS at the required concentration (approximately 1.5×10^8^ bacteria/mL). Phenol Red (Sigma, P0290-100 mL, 0.5% phenol red solution diluted 0.1% in the injected solution) and fluorescent dye (Dextran, tetramethylrhodamine, 10,000 MW, anionic, D-1868 Invitrogen, 30 µM in the injected solution) were added to the suspension to facilitate monitoring of the injection process. The GAB zebrafish line was used for all trials, which were carried out in fish water at 28° C. Infections with the *S. aureus* NSA1385, Δ*rosa*, and Δ*rosa* + pTSS::*ssbP* strains were achieved by injecting 1 nL of bacterial suspensions into the duct of Cuvier of 50 hours post-fertilization (hpf) embryos that had been dechorionated and anesthetized with tricaine (0.3 mg/mL). Each embryo was injected individually and placed into a separate well of a 96-well microtiter plate. To confirm the bacterial load per injection, the same volume was injected into 200 µL of PBS, and the number of CFU was determined using TSB agar plates. Survival kinetics were assessed visually by counting the number of embryos with an absence of heartbeat. Infections with *S. aureus* USA300 were performed using the same general protocol but with some modifications. Injections were carried out in the hindbrain ventricle of 30 hpf embryos, a site chosen to ensure the close proximity of all injected components and enable localized effects. The control buffer or recombinant SSBP protein was injected at the same site 30 minutes after the initial bacterial injection. The protein was administered at a maximum concentration of 0.13 µM per embryo to avoid precipitation. The volume injected was also limited by technical constraints, as the hindbrain ventricle cannot accommodate larger amounts.

### Ethical Statement

All animal experiments were carried out at the University of Montpellier in accordance with European Union recommendations for the care and use of laboratory animals. The study was authorized by the *Direction Sanitaire et Vétérinaire de l’Hérault* and *Comité d’Ethique pour l’Expérimentation Animale* under reference CEEALR-B4-172-37 and APAFIS#5737-2016061511212601 v3. Adult zebrafish were not euthanized for this study and were bred in accordance with the international standards outlined in the EU Animal Protection Directive 2010/63/EU. All experiments were performed prior to the embryos’ free-feeding stage and, as such, were not classified as animal experimentation under the EU Animal Protection Directive 2010/63/EU. For survival analysis, the presence or absence of a heartbeat was used as the clinical criterion. Once survival monitoring was completed, the plates were frozen immediately and stored at −20°C for 48 h to ensure embryo death. Subsequently, they were autoclaved along with other bacterial-contaminated waste.

### RNA-Sequencing Analysis

RNA-sequencing was performed using the RNeasy Plus Mini Kit (Qiagen) following the manufacturer’s instructions with minor modifications. Total RNA was extracted from *S. aureus*-infected RAW 264.7 macrophages. RAW 264.7 cells were seeded in a 6-well plate at a density of 5 × 10⁵ cells/mL. After 24 h, infection was carried out as previously described, resulting in approximately 4 × 10⁶ infected cells. At the designated time point (T5h), cells were detached using a cell scraper and collected into a 50 mL tube. The cell suspension was centrifuged at 4,000 g for 10 min, and the supernatant was carefully removed by pipetting. According to the manufacturer’s recommendations, the volume required to dissolve the pellet was calculated, and RNAprotect Cell Reagent (Qiagen), lysostaphin (50 µg/mL), and 0.1% Triton X-100 were added to the sample. The mixture was vortexed and further homogenized using a needle to maximize RNA yield and prevent clogging of the RNeasy spin column. Isolated RNA samples were resuspended in nuclease-free water to eliminate potential contaminants that could interfere with downstream analyses. RNA quality and quantity were assessed using a NanoDrop One spectrophotometer. RNA samples were sent to Novogene (Cambridge, UK) for sequencing. Novogene prepared RNA sequencing libraries on the Illumina NovaSeq 6000 platform. Each sample contained 2.5 µg of total RNA, which was converted into cDNA libraries. Sequencing parameters were set to generate 100 million paired-end reads per sample, with each read measuring 150 bp.

### Enzyme-Linked Immunosorbent Assay (ELISA)

The levels of IL-6, TNF-α, and IL-10 cytokines in the supernatants of *S. aureus*-infected RAW 264.7 macrophages were measured using ELISA kits (Proteintech, Wuhan, China) according to the manufacturer’s instructions. Supernatants were collected at designated time points post-infection and clarified by centrifugation to remove cellular debris. Each experimental sample was analyzed in triplicate, and cytokine concentrations were quantified by comparison to standard curves generated from serial dilutions of known cytokine standards provided in the kit. The results were expressed as mean cytokine concentrations (pg/mL) ± standard deviation (SD) from three independent experiments.

### Statistics and Bioinformatics

The statistical significance of differences between groups was estimated using the GraphPad software package Prism 9.5.1, as stated in the figure legends. For the bioinformatics analysis, gene ontology (GO) classification was conducted using the ShinyGO platform to identify enriched categories (Ge *et al*., 2020). Additionally, REACTOME pathway enrichment analysis was performed with the mouse genome as the reference background. Pathways with an FDR-adjusted p-value below 0.05 were considered statistically significant and extracted for further analysis.

### Data and Resource Availability

All data generated or analyzed during this study are included in the published article and its online supplementary files. No applicable resources were generated or analyzed specifically for this study.

## RESULTS

### Transcriptomic analysis highlights ROSA-like prophage gene overexpression in *S. aureus* NSA1385

We conducted a transcriptomic analysis to compare the gene expression profiles of *S. aureus* NSA1385 and the reference *S. aureus* Newman strains under standard growth conditions. The *S. aureus* Newman strain harbors a prophage at the same genomic locus as the ROSA-like prophage in NSA1385, with a sequence homology of 95% (Bae *et al*., 2006; Messad *et al*., 2015). Given that the attenuated virulence profile of NSA1385 is strongly associated with the presence of the ROSA-like prophage, we focused on identifying overexpressed genes within this prophage that could explain the observed phenotype. Our microarray analysis revealed that three ORFs within the ROSA-like prophage, ORF35, ORF22, and ORF15, were significantly overexpressed in NSA1385 compared to Newman **(Table 1)**. ORF35 and ORF22 showed 10-fold and 6-fold increases, respectively, while ORF15 displayed a remarkable 23-fold increase in expression under standard growth conditions. These findings highlight the potential role of the ROSA-like prophage in modulating the behavior of NSA1385.

**Table 1.** Differential expression of ROSA-like prophage genes in NSA1385 compared to the Newman strain using a microarray approach.

| Locus | Description | Fold change NSA1385 vs Newman |
| --- | --- | --- |
| UniRef100_A5IRK3 | *orf4* | 0.12 |
| UniRef100_A6TYD4 | *orf55* | / |
| UniRef100_A9CR75 | *orf22* | 6.35 |
| UniRef100_A9CR68 | *orf17* | / |
| UniRef100_A9CR94 | *orf35* | 10.35 |
| UniRef100_A9CRC4 | *orf56* | 0.21 |
| UniRef100_A9CRD3 | *orf62* | 0.31 |
| UniRef100_B2ZYV1 | *orf13* | / |
| UniRef100_B2ZYW4 | *orf26* | 0.15 |
| UniRef100_B2ZYY5 | *orf47* | 0.29 |
| UniRef100_B2ZZ04 | *orf66* | 0.24 |
| UniRef100_P37455 | *orf15 (ssbP)* | 23.60 |
| UniRef100_Q4ZA35 | *orf38* | 0.25 |
| UniRef100_Q4ZAE7 | *orf19* | / |
| UniRef100_Q4ZAE9 | *orf40* | / |
| UniRef100_Q4ZAF0 | *orf53* | / |
| UniRef100_Q4ZAF8 | *orf57* | 0.21 |
| UniRef100_Q4ZAH3 | *orf48* | 0.19 |
| UniRef100_Q4ZAH4 | *orf47* | 0.09 |
| UniRef100_Q4ZAZ9 | *orf58* | / |
| UniRef100_Q4ZBU5 | *orf78* | / |
| UniRef100_Q4ZBZ1 | *orf132* | / |
| UniRef100_Q4ZDP3 | *orf89* | / |
| UniRef100_Q9MBT2 | *orf7* | / |

### ORF15 codes for a Single-Stranded DNA-binding protein

Among the three overexpressed ORFs identified in the ROSA-like prophage, genomic analyses using the National Center for Biotechnology Information (NCBI) databases revealed distinct characteristics. ORF35 and ORF22 were predicted to encode hypothetical proteins with no currently identified functions. In contrast, ORF15 exhibited significant homology to single-stranded DNA-binding proteins, a well-characterized class of proteins known for their ability to bind ssDNA with high affinity, irrespective of the DNA sequence (Guo and Malik, 2022). Based on these findings, we designated ORF15 as SSBP. Furthermore, among the 278 protein homologs retrieved from the UniProt database, 52 were selected for detailed analysis, including those from bacteriophages, bacteria, and mitochondria of protozoa, plants, and animals. Notable examples include the human mitochondrial single-stranded DNA-binding protein and the chloroplastic OSB2 protein from *Arabidopsis thaliana* (**Sup.** Fig. 1). SSBP demonstrated 100% identity to the *Staphylococcus* phage phiMR25 protein (GenBank accession n°YP001949814) within the *Caudoviricetes* family, and 99% sequence similarity to *S. aureus* SSB1. In contrast, SSBP shared less than 55% sequence similarity with non-phage homologues (**Sup.** Fig. 1). The characterized SSBP monomer comprises 141 amino acids with a molecular mass of 19 kDa. Sequence analysis revealed a conserved C-terminal tail (IEDLPF), analogous to the DDDIPF motif in *E. coli* homologue, known for mediating protein–protein interactions (Antony *et al*., 2013) (**Sup Fig. 2**). Additionally, SSBP features four canonical domains of single-stranded DNA-binding proteins: the oligonucleotide/oligosaccharide/oligopeptide-binding (OB) fold, K Homology (KH) domain, RNA Recognition Motif (RRM), and Whirly domain (Marceau, 2012). In order to investigate the functionality of SSBP, the protein was cloned, overexpressed, and purified. Binding affinity assays were conducted using Electrophoretic Mobility Shift Assays (EMSAs) to assess interactions with ssDNA and dsDNA. Increasing concentrations of recombinant SSBP (0.5 to 2.5 µM) were incubated with a constant concentration of an 80-mer oligonucleotide probe labeled with fluorescent dye DY682. The results demonstrated that SSBP specifically binds ssDNA in a dose-dependent manner, while no binding was observed with dsDNA (**Fig. 1A**). These findings confirm that SSBP is a high-affinity ssDNA-binding protein, consistent with its predicted function.

**Figure 1.**
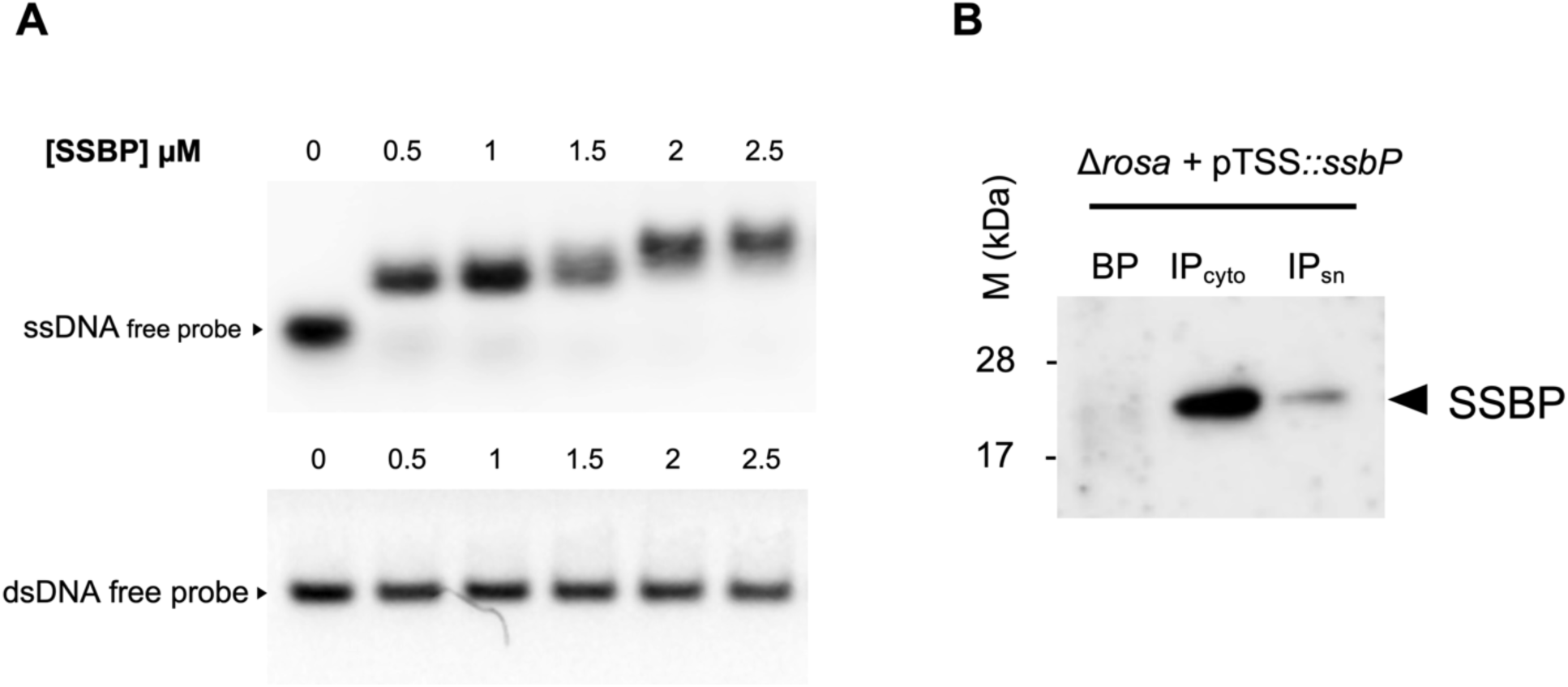
SSBP is a secreted Single-Stranded DNA-binding protein. **(A)** Electrophoretic mobility shift assay (EMSA) of SSBP to ssDNA (upper panel) and dsDNA (lower panel) probes. Purified recombinant N-terminally 6x-His-tagged SSBP was incubated at increasing concentrations (0.5 to 2.5 µM) with a constant concentration of an 80-mer oligonucleotide probe (100 fmol)labeled with DY682. **(B)** Supernatants from Δ*rosa* + pTSS::*ssbP*-Spot cultures were filtered and subjected to immunoprecipitation using anti-Spot magnetic beads. Bacterial pellets were resuspended and lysed in PBS containing lysostaphin, a protease inhibitor cocktail, and DNase I. SDS-PAGE was performed to analyze bacterial pellets (BP) and immunoprecipitated proteins from the supernatant (IPsn) and cytosol (IPcyto). Proteins were transferred to PVDF membranes and analyzed by Western blotting using a monoclonal mouse anti-Spot antibody (Chromotek, Germany) as the primary antibody and an HRP-conjugated donkey anti-mouse antibody (Jackson ImmunoResearch) as the secondary antibody. Data are representative of three independent experiments.

### SSBP has no impact on *S. aureus* growth and is secreted *in vitro*

To mimic the overexpression levels of *ssbP* observed in the transcriptomic results, we constructed the pTSS::*ssbP* plasmid to overexpress the SSBP protein. Given the compact genomic organization of the prophage locus, where *ssbP* is embedded among closely spaced co-transcribed genes, targeted deletion of *ssbP* was not feasible without risking polar effects. We therefore used the prophage-excised strain ΔROSA to assess the specific contribution of *ssbP* via genetic complementation. The NSA1385 prophage-excised strain (ΔROSA) was transformed with either the empty vector plasmid or the plasmid overexpressing SSBP, yielding the ΔROSA + pTSS and ΔROSA + pTSS::*ssbP S. aureus* strains, respectively. Growth analysis revealed no significant differences in the exponential growth phase among these strains **(Sup. Fig. 3)**. These results suggest that neither the presence of the ROSA-like prophage nor the overexpression of SSBP affects the *in vitro* growth of *S. aureus*.

To assess whether SSBP could be secreted, we focused on the overexpression system, which not only reflects the high levels detected in transcriptomic datasets but also facilitates protein detection. Because no anti-SSBP antibody was available, using a tagged and overexpressed version ensured reliable monitoring of protein secretion and minimized the risk of missing secretion due to low endogenous expression levels. We therefore used immunoprecipitation with anti-Spot magnetic beads to analyze the supernatants and cytosolic fractions of *S. aureus* cultures for the presence of SSBP-Spot. A strong SSBP-Spot signal was detected in the cytosolic fraction, confirming its intracellular production **(Fig. 1B)**. Interestingly, a weaker yet discernible signal was observed in the supernatant, suggesting that the protein is partially secreted despite lacking a classical secretion signal sequence **(Fig. 1B)**. These findings demonstrate that SSBP is secreted, at least partially, during *in vitro* growth, supporting its hypothesized role as a secreted factor potentially contributing to intracellular infection.

### SSBP is involved in the decreased survival of the colonizing *S. aureus* strain during macrophage infection

To further elucidate the interaction between *S. aureus* NSA1385 and the immune system, we examined the role of the ROSA-like prophage and SSBP during macrophage infection. Murine RAW 264.7 macrophages and primary human THP-1 macrophages were used to phagocytose the *S. aureus* strains, and intracellular viable bacteria were quantified at 5 and 24 h post-Gentamicin treatment (pGt). All tested strains exhibited similar CFU at T0, indicating that neither the ROSA-like prophage nor SSBP influenced phagocytosis **(Fig. 2A and 2D)**.

**Figure 2.**
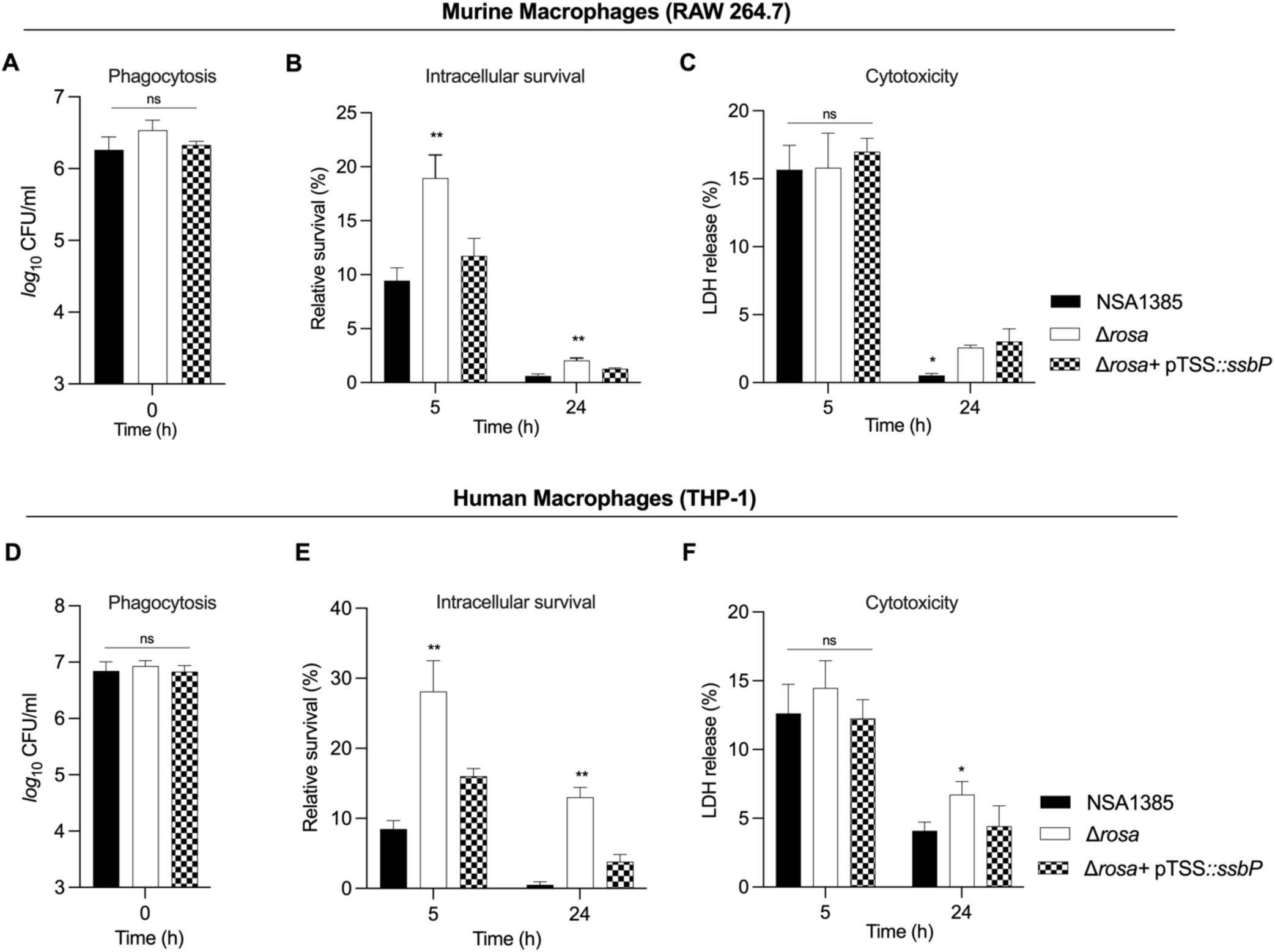
SSBP expression reduces survival in infected macrophages. *S. aureus* NSA1385, Δ*rosa*, and Δ*rosa* + pTSS::*ssbP* bacteria were used to infect RAW 264.7 (A-C) and THP-1 (D-F) macrophages. (A and D) Phagocytosis of *S. aureus* strains by macrophages was evaluated by infecting cells at a MOI of 20 for 1 h at 37 °C. After infection, cells were incubated with gentamicin for 30 minutes to eliminate extracellular bacteria, followed by incubation in complete media supplemented with lysostaphin to kill extracellular bacteria released from lysed macrophages during the subsequent. Cells were then lysed in 0.1% Triton X-100. (B and E) Surviving bacteria in macrophages lysates were evaluated by counting CFUs at 5 and 24 h after post-lysostaphin/gentamicin treatment. Bacterial survival rate was estimated by dividing the number of bacterial colonies at the post-gentamicin time point by the number of colonies at T0 and multiplying by 100. (C and F) LDH release was measured using the CyQUANT assay kit. RAW 264.7 and THP-1 macrophages were infected at a MOI of 20 for 5 and 24 h. For these experiments, cells were seeded in a 96-well plate, and extracellular bacteria were not eliminated to allow for LDH release measurement. Data represents the mean + SD of five independent experiments. Statistical significance was determined using two-way ANOVA (**p < 0.01; *p < 0.05; ns, not significant).

However, a significant reduction in intracellular survival was observed for the NSA1385 strain compared to Δ*rosa* at both 5 and 24 h pGt **(Fig. 2A and 2D)**. This suggests that excision of the ROSA-like prophage restored the ability of *S. aureus* to survive within macrophages. Interestingly, the strain overexpressing *ssbP* exhibited a similarly reduced intracellular survival rate, comparable to the NSA1385 strain harboring the ROSA-like prophage **(Fig. 2B and 2E)**. This finding highlights the role of SSBP in decreasing the intracellular survival of the colonizing strain NSA1385 within both murine and human macrophages.

To determine whether the reduced intracellular survival of NSA1385 and the *ssbP*-overexpressing strains was related to increased macrophage cytotoxicity, we performed LDH assays. This assay measures plasma membrane damage as an indicator of necrotic cell death by detecting LDH enzyme release into the culture medium. Our results revealed no significant differences in cytotoxicity among the three strains, indicating that the observed differences in survival were independent of macrophage cell death **(Fig. 2C and 2F).**

### SSBP is not involved in the intracellular survival of *S. aureus* NSA1385 in non-phagocytic cells

Next, we sought to investigate whether SSBP was involved in the intracellular survival of *S. aureus* within different cell types, particularly non-phagocytic cells, as previous studies demonstrated the role of the ROSA-like prophage in osteoblasts for the NSA1385 strain (Rasigade *et al*., 2016). Human skin keratinocytes (HaCaT), predominant in the epidermis and critical for wound healing (Secor *et al*., 2011), and MC3T3-E1 murine osteoblasts were used to evaluate the impact of SSBP on *S. aureus* internalization and persistence.

Our results revealed that the NSA1385 strain exhibited significantly higher internalization in keratinocytes at T0 compared to the other two strains, highlighting the ROSA-like phage role in enhancing the internalization ability of the colonizing strain **(Fig. 3A).** However, this higher internalization was not observed in osteoblasts, where all three strains exhibited similar CFU counts at T0 **(Fig. 3D)**. Over time, the intracellular load of NSA1385 was significantly lower compared to the Δ*rosa* and *ssbP*-overexpressing strains, which demonstrated higher persistence and intracellular survival in both osteoblasts and keratinocytes **(Fig. 3B and 3E)**. These findings confirm that the ROSA-like phage contributes to reduced survival of the colonizing strain in osteoblasts and keratinocytes, though independent of SSBP.

**Figure 3.**
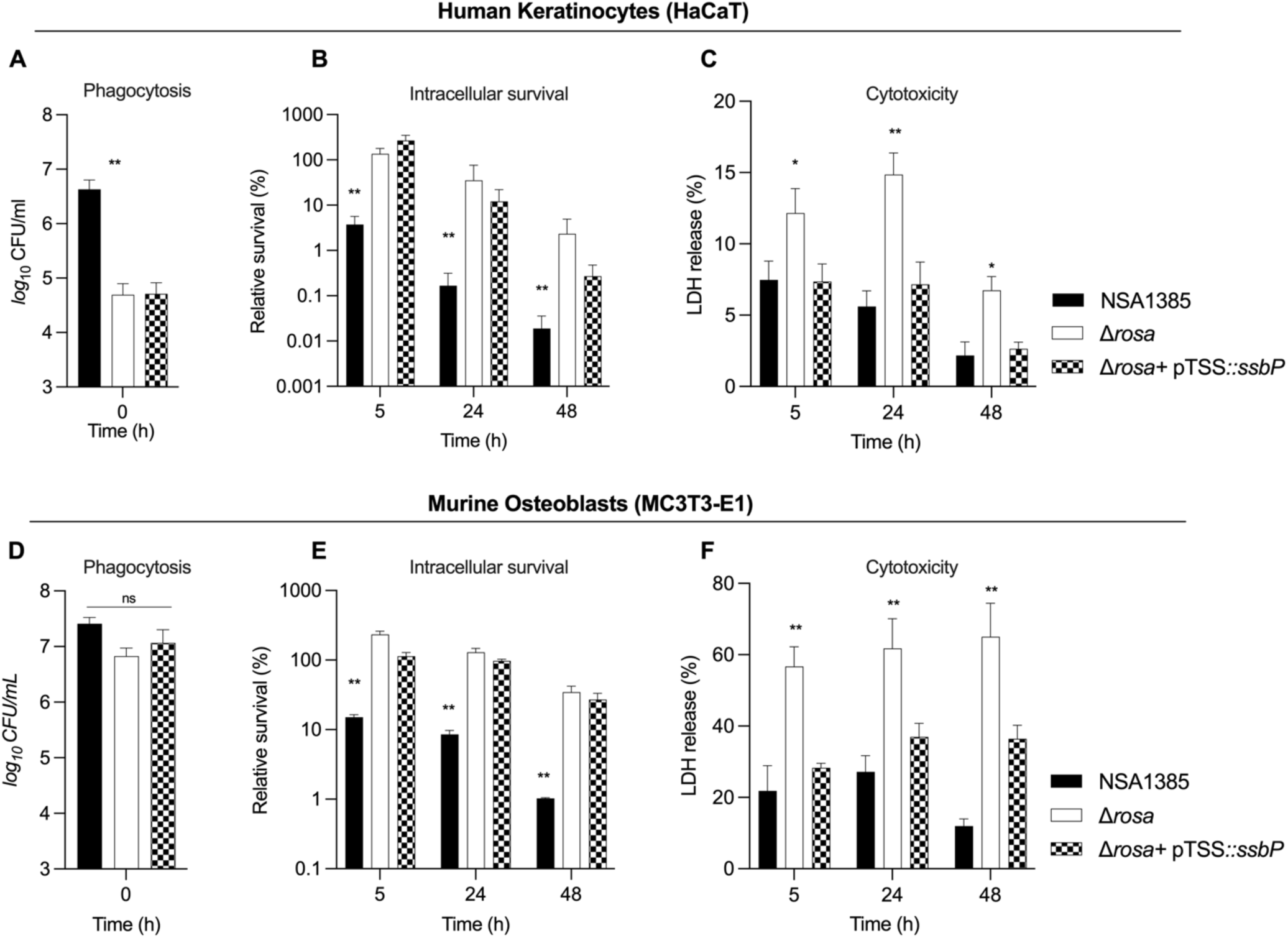
Effect of SSBP expression in infected keratinocytes and osteoblasts. *S. aureus* NSA1385, Δ*rosa*, and Δ*rosa* + pTSS::*ssbP* strains were used to infect HaCaT keratinocytes (A-C) and MC3T3-E1 osteoblasts (D-F). Cells were infected at a MOI of 100 for 1h30 at 37 °C, followed by a 30 minutes incubation with gentamicin to eliminate extracellular bacteria. Cells were then incubated in complete media supplemented with lysostaphin to kill extracellular bacteria released from lysed cells during the subsequent incubation period. At the indicated time points, cells were lysed in 0.1% Triton X-100. Phagocytosis (T0) and intracellular survival (5, 24, and 48 h) were evaluated by determining CFUs in cell lysates. The bacterial survival rate was calculated as the percentage of CFUs at each post-gentamicin time point relative to CFUs at T0. Cytotoxicity was assessed by measuring LDH release using the CyQUANT assay kit. For these experiments, cells were seeded in 96-well plates and infected at a MOI of 100 for 5, 24, and 48 h, without eliminating extracellular bacteria to allow for LDH quantification. Data are presented as the mean ± SD of five independent experiments. Statistical significance was determined using two-way ANOVA (**p < 0.01; *p < 0.05; ns, not significant).

Furthermore, infection with the Δ*rosa* strain increased LDH release in both keratinocytes and osteoblasts over time, indicating a correlation between high intracellular survival and increased cytotoxicity **(Fig. 3C and 3F)**. Conversely, cells infected with NSA1385 and Δ*rosa* + pTSS::*ssbP* strains exhibited significantly reduced LDH release over time, consistent with lower intracellular survival **(Fig. 3C and F).**

Overall, these results demonstrate that the ROSA-like prophage reduces intracellular survival in both keratinocytes and osteoblasts, independently of SSBP.

### RNA-sequencing analysis upon macrophage infection revealed SSBP-dependent up-regulation of immune response pathways

To investigate the mechanisms underlying the reduced survival of the ΔROSA + pTSS::*ssbP* strain during macrophage infection, we performed dual RNA-Seq analysis to assess both the bacterial and host transcriptional responses. Gene expression profiles were compared between macrophages infected with the ΔROSA + pTSS and those infected with the ΔROSA + pTSS::*ssbP S. aureus* strains. We first analyzed the bacterial transcriptome recovered from infected macrophages to determine whether SSBP expression altered the expression of bacterial genes, including virulence factors. The comparison revealed no significant differences between the two strains, suggesting that SSBP does not impact the bacterial transcriptional program during intracellular infection. However, the host transcriptome showed substantial differences in response to SSBP. Our analysis identified a total of 1,211 differentially expressed genes (DEGs) exhibiting at least a twofold change with a *P*-value < 0.05. Of these, 764 genes were upregulated, while 447 were downregulated in response to SSBP overexpression **(Sup. Table S3)**. To investigate the cellular pathways associated with the identified DEGs, gene set enrichment analysis was carried out using the ShinyGO tool (Ge *et al*., 2020). Reactome pathway enrichment analysis revealed that up-regulated DEGs were predominantly associated with the immune system, cytokine signaling, and signal transduction pathways **(Fig. 4A)**. For example, pathways such as interleukin signaling and TNFR2 non-canonical NF-kB signaling were highlighted, emphasizing their roles in immune cell activation and inflammatory response to *S. aureus* infection. The most overrepresented pathways included interleukin signaling, TNFR2 non-canonical NF-kB signaling, and CLEC7A Dectin-1 signaling. In addition, the most overrepresented biological process terms included response to stress, immune response, response to cytokine stimulus, and defense response **(Fig. 4B)**. For molecular function, terms including cytokine activity, signaling receptor binding, and receptor regulator activity are the most significant **(Fig. 4C)**. However, a similar analysis was conducted for the downregulated genes, and no significantly enriched pathways could be identified for this gene set **(Sup. Table S3).** This absence of enriched pathways may suggest that the downregulated genes are either more diverse in their functional roles. Therefore, our results demonstrate that SSBP selectively modulates the host immune response without altering bacterial gene expression, particularly enhancing interleukin signaling and other macrophage activation pathways.

**Figure 4.**
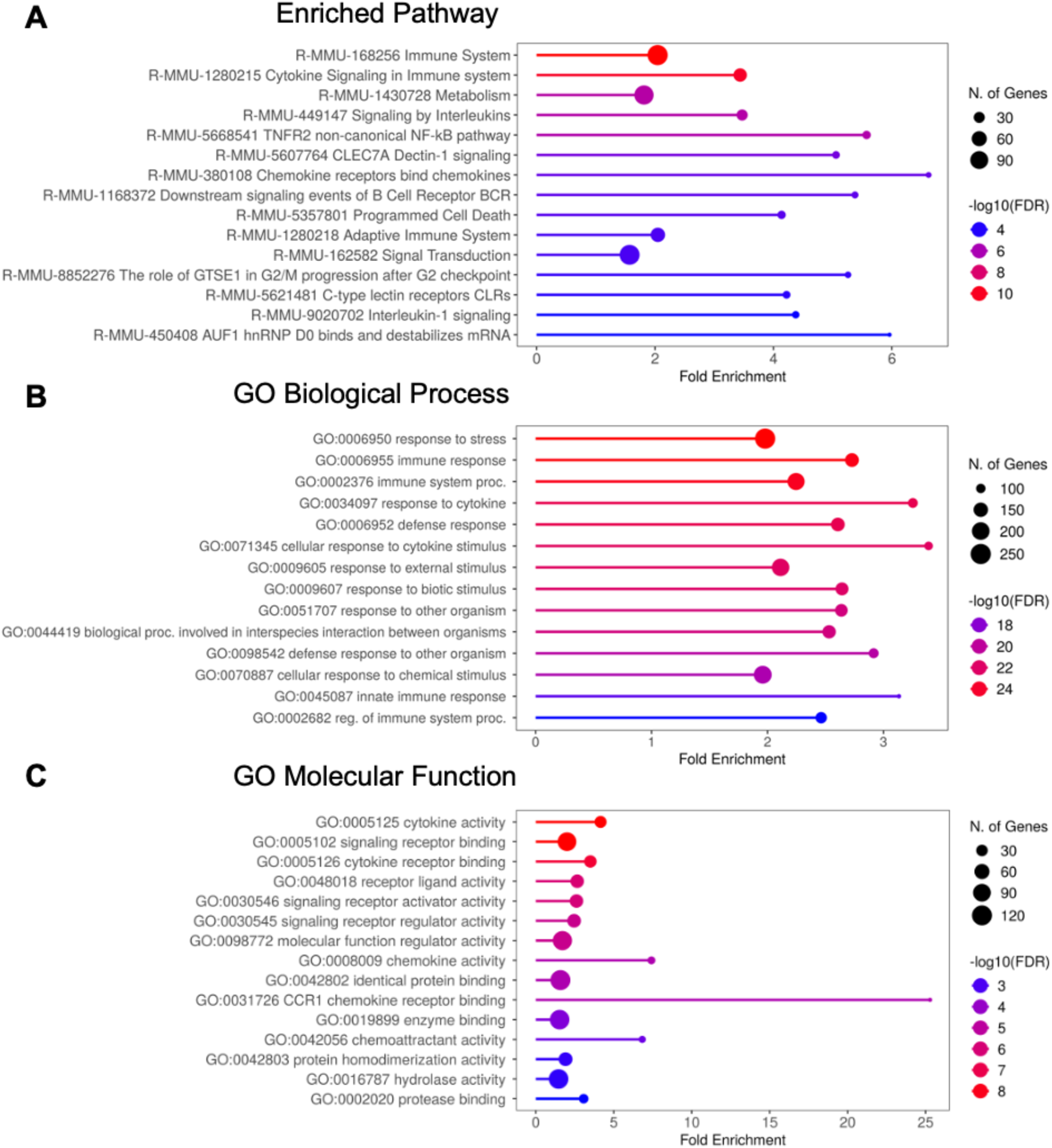
SSBP-correlated transcripts in macrophages infected with Δ*rosa* + pTSS or Δ*rosa* + pTSS::*ssbP S. aureus* strains and enrichment analysis using ShinyGO. (A), (B), and (C) represent the significant roles of up-regulated DEGs in enriched pathways, biological processes, and molecular functions, respectively. The terms are ranked by fold enrichment, with colors representing −log10(FDR) values derived from nominal p-values calculated through the hypergeometric test. Circle size reflects the number of genes associated with each term.

### SSBP modulates cytokine response during infection

The innate immune system, as the first line of defense, orchestrates a cascade of pro-inflammatory signals and cytokine release during bacterial infection (Fournier and Philpott, 2005). Based on our RNA-seq analysis, we investigated the immunomodulatory effects of SSBP and examined its impact on key cytokines involved in macrophage-mediated inflammatory responses. Using ELISA assays, we quantified the secretion levels of IL-6, TNF-α, and IL-10 by macrophages infected with the colonizing strain NSA1385, the ROSA-like prophage-deficient strain Δ*rosa*, and the Δ*rosa* strain overexpressing SSBP (Δ*rosa* + pTSS::*ssbP*). At T0 (immediately post-gentamycin treatment), cytokine levels were comparable to those of non-infected macrophages, indicating baseline cytokine production. However, by T5h (once infection was established), cytokine production increased markedly across all conditions **(Fig. 5)**. Specifically, macrophages infected with NSA1385 or Δ*rosa* exhibited similar IL-6 secretion (∼500 pg/mL), whereas Δ*rosa* + pTSS::*ssbP* triggered a significantly elevated IL-6 response (∼1500 pg/mL) **(Fig. 5A)**, highlighting the pronounced pro-inflammatory effect of SSBP overexpression. Similarly, TNF-α levels increased in all strains, with a marginally higher production observed in macrophages infected with the SSBP-overexpressing strain **(Fig. 5B)**. In contrast, IL-10, an anti-inflammatory cytokine, was markedly reduced in macrophages infected with NSA1385 or Δ*rosa* + pTSS::*ssbP*, with levels dropping below 100 pg/mL, compared to the relatively higher IL-10 secretion (∼150 pg/mL) observed in macrophages infected with Δ*rosa* **(Fig. 5C)**. These results suggest that SSBP enhances the inflammatory response, particularly by elevating IL-6 levels, which may facilitate the clearance of macrophages. However, in the presence of SSBP, this increase in IL-10 is inhibited. The combination of heightened pro-inflammatory and reduced anti-inflammatory cytokine levels likely explains the more rapid elimination of the colonizing strain from macrophages, highlighting the significant role of SSBP in modulating immune responses during infection.

**Figure 5.**
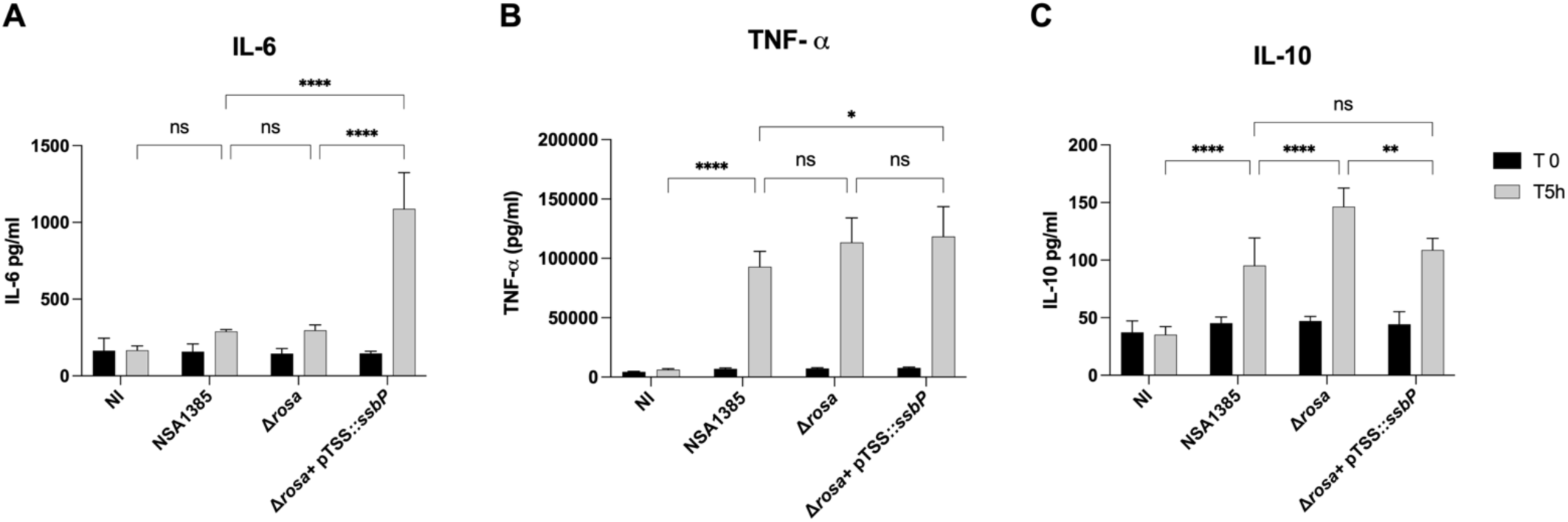
Effect of *S. aureus* NSA1385, Δ*rosa*, and Δ*rosa* + pTSS::*ssbP* infections on cytokines production in RAW 264.7 macrophages. Cytokines levels in the supernatants of macrophage cultures were collected at 0 and 5 h pGt and measured using ELISA assays. “NI” corresponds to non-infected macrophages. Data are presented as the mean ± SD of three independent experiments. Statistical significance was determined using two-way ANOVA (ns, not significant; *p < 0.05; **p < 0.01; ****p < 0.0001).

### Recombinant SSBP protein reduces *S. aureus* intramacrophagic survival across diverse clinical strains

Given the observed effect of the SSBP-overexpressing strain on modulating macrophage responses and reducing bacterial survival, we sought to investigate whether the recombinant SSBP protein itself could directly influence the intramacrophagic survival of various *S. aureus* clinical strains during macrophage infection. This included the Δ*rosa* strain; SA564, a clinical isolate belonging to clonal complex 5; NSA739, isolated from a grade 3 DFU; and USA300, a highly virulent MRSA strain. We infected RAW 264.7 macrophages with the Δ*rosa*, SA564, NSA739, and USA300 strains to investigate their intracellular survival. All strains exhibited similar levels of phagocytosis at T0 **(Fig. 6A**).

**Figure 6.**
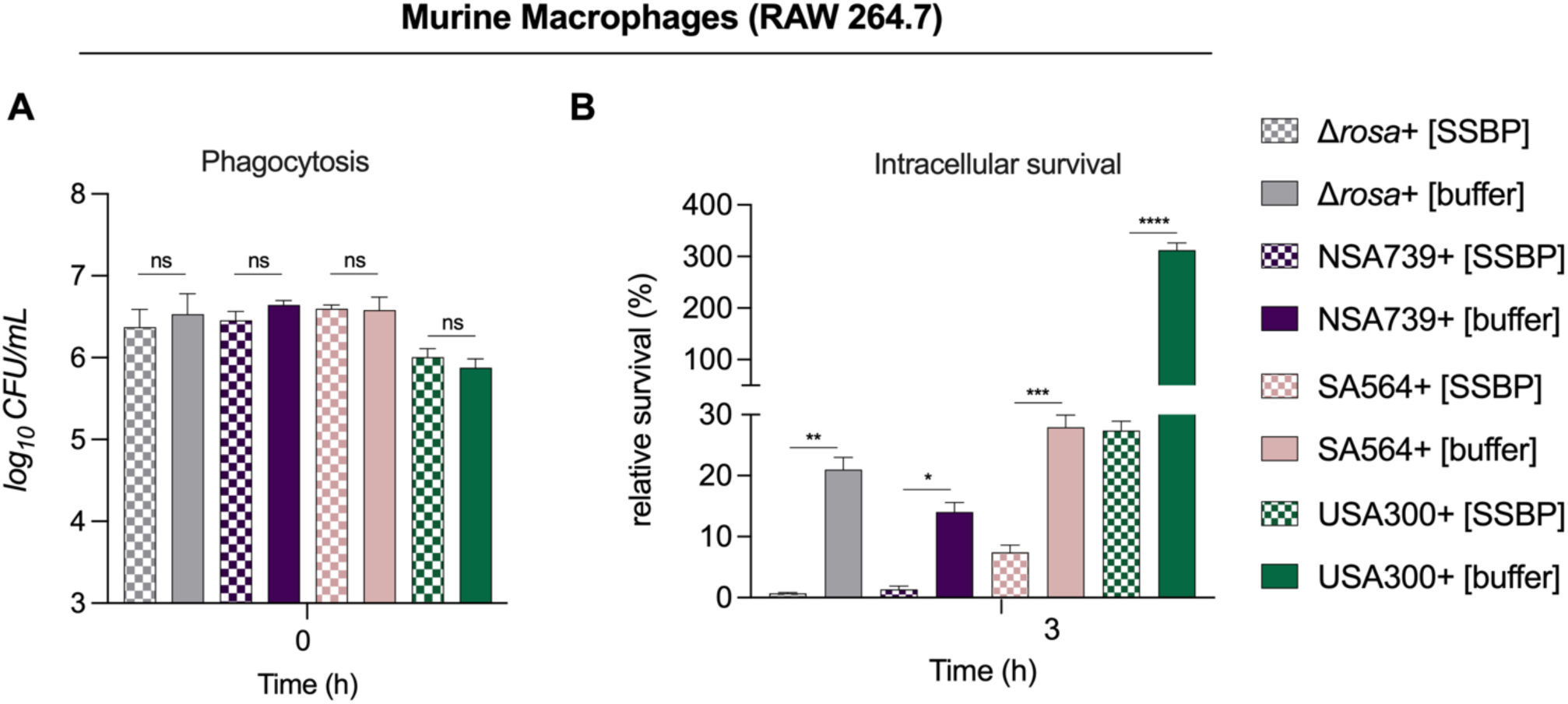
Pure recombinant SSBP protein reduces *S. aureus* intramacrophagic survival. Δ*rosa*, NSA739, SA564, USA300 strains were used to infect RAW 264.7 macrophages **(A)** Phagocytosis: Macrophages were infected at a multiplicity of infection (MOI) of 20, without or with the addition of 3 µM recombinant SSBP for 1 h at 37°C. After infection, cells were incubated with gentamicin for 30 minutes to eliminate extracellular bacteria, followed by incubation in complete media supplemented with recombinant SSBP and lysostaphin to kill any extracellular bacteria released during the subsequent incubation. Cells were then lysed in 0.1% Triton X-100 for CFU enumeration. **(B)** Intracellular survival: Macrophages were lysed, and surviving bacteria were quantified by counting CFUs in cell lysates 3 h after lysostaphin/gentamicin treatment. The bacterial survival rate was calculated as the percentage of CFUs at the post-gentamicin time point relative to CFUs at T0. Data are presented as the mean ± SD of five independent experiments. Statistical significance was determined using two-way ANOVA (*p < 0.05; **p < 0.01; ***p < 0.001; ****p < 0.0001; ns, not significant).

Remarkably, the addition of recombinant SSBP significantly reduced the intracellular survival rates across all tested strains at 3 h pGt compared to the control condition (buffer only). Specifically, the Δ*rosa* strain showed a 90% reduction in survival, NSA739 exhibited an 80% reduction, and USA300 demonstrated a striking 90% reduction in survival **(Fig. 6B).** Notably, USA300 actively divides within macrophages, achieving a survival rate of 300% in the absence of SSBP. This highlights the potent effect of SSBP in countering the aggressive intracellular proliferation of this highly virulent strain.

A concentration of 3 µM was determined to be the minimal effective dose to achieve this reduction, and further time-course experiments indicated that 3 h of incubation was sufficient to observe significant effects (**Fig. 6B**). Interestingly, when macrophages were pre-treated with SSBP for 2 h prior to infection, no changes in phagocytosis or intracellular survival were observed, indicating that SSBP does not directly activate macrophages but rather acts during the intracellular survival phase, in accordance with IL-6 production.

These findings demonstrate that recombinant SSBP protein effectively reduces *S. aureus* survival across diverse strains, including the highly virulent MRSA USA300, and suggest its potential as a therapeutic agent targeting intramacrophagic persistence of *S. aureus*.

### SSBP Attenuates Virulence in Zebrafish Infected Embryos

Given the demonstrated role of SSBP in reducing the intracellular survival of *S. aureus* within macrophages and its impact on pro-inflammatory cytokine production, we sought to evaluate its effect in an *in vivo* infection model. To this end, we employed the zebrafish embryo model, a well-established tool for evaluating bacterial virulence by monitoring the survival of infected embryos. First, we injected *S. aureus* NSA1385 and its isogenic Δ*rosa* derivative strains into the duct of Cuvier of zebrafish embryos at 50 hpf, which induces a systemic infection. Analysis of survival at 80 h post-infection (hpi) revealed that NSA1385 injection resulted in approximately 70% survival, while the Δ*rosa* strain showed a significantly lower survival rate of 20% **(Fig. 7A)**. This confirms our previous findings demonstrating the attenuation of virulence in the colonizing NSA1385 strain (Messad *et al*., 2015). Interestingly, embryos infected with the Δ*rosa* + pTSS::*ssbP* strain recapitulated a higher survival rate similar of NSA1385.

**Figure 7.**
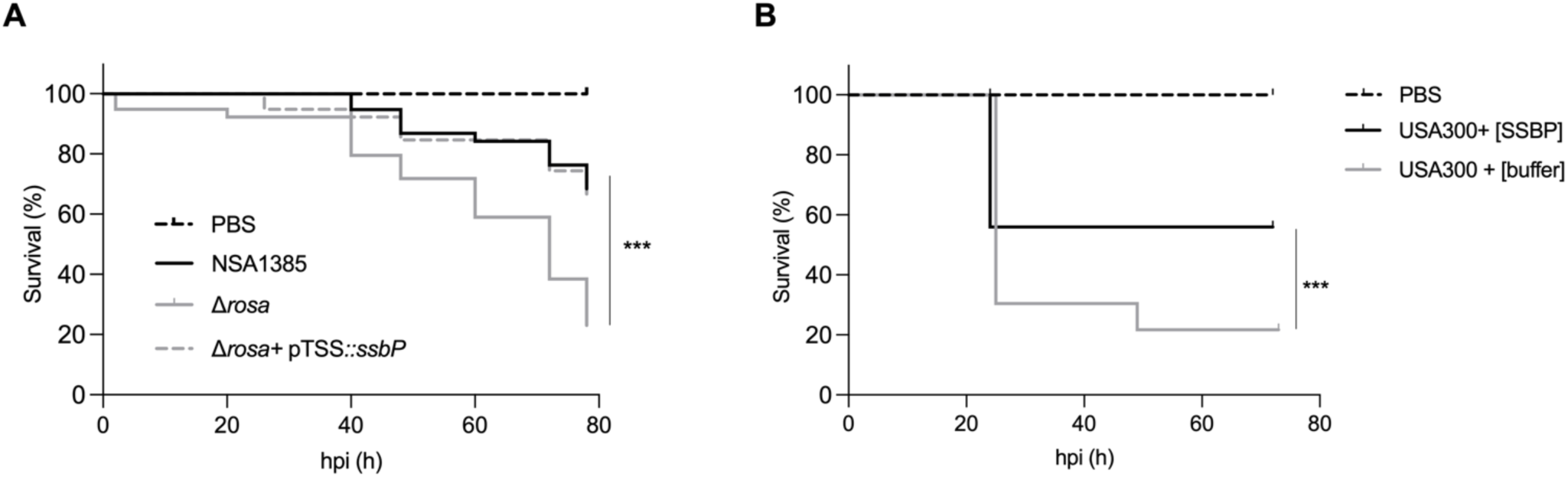
SSBP attenuates virulence in infected zebrafish embryos. (A) Kaplan-Meier representation of the survival of zebrafish embryos infected via injection into the duct of Cuvier with NSA1385, Δ*rosa*, and Δ*rosa*+ pTSS::*ssbP* at 3.10^3^ CFU/nL grown in the exponential phase, or PBS (negative control) The proportion of surviving embryos (n = 50 per group, representative of three independent experiments) is shown. (B) Kaplan-Meier representation of the survival of zebrafish embryos infected via injection into the hindbrain ventricle with USA300 at 2.10^3^ CFU/nL, followed 30 minutes later by injection of recombinant SSBP protein (0.13 µM) or buffer control at the same injection site, or PBS (negative control). The proportion of surviving embryos (n = 30 per group, representative of three independent experiments) is shown. Statistically significant differences in survival between the strains are indicated with asterisks: *** p < 0.001 (log-rank test). hpi, hours post-infection.

Next, building on the observed reduction in bacterial load in macrophages infected with the highly virulent USA300 strain following recombinant SSBP treatment, we sought to evaluate its *in vivo* impact. Zebrafish embryos were injected with *S. aureus* USA300 and administered 30 minutes post-infection with either a buffer control or recombinant SSBP protein at a concentration of 0.13 µM/embryo, which represents the maximum concentration of pure recombinant protein that could be tested without risking protein precipitation. Survival analysis over 80 hpi revealed a striking difference between the two conditions **(Fig. 7B).** Embryos infected with USA300 treated with buffer control showed a rapid decline in survival, with less than 20% of embryos surviving by the end of the experiment. In contrast, embryos treated 30 minutes post-infection with recombinant SSBP following infection with USA300 exhibited significantly enhanced survival, with approximately 70% of embryos remaining alive at 80 hpi. These results are representative of multiple experiments in which the concentration of USA300 CFUs was varied from 2500 to 8000 CFU per embryo. Survival rates remained consistent across all tested conditions, demonstrating the efficacy of SSBP even at very high bacterial loads. Notably, lower bacterial doses, such as 2500 CFU, exhibited a similar survival-enhancing effect, indicating that the impact of SSBP is independent of bacterial load. Therefore, the observed *in vivo* attenuation of USA300 virulence aligns with the *in vitro* macrophage survival assays, reinforcing the role of recombinant SSBP protein, in reducing *S. aureus* infection.

## DISCUSSION

Normally, pathogenic bacteria, including *S. aureus*, exploit immunosuppressive mechanisms to promote their survival and persistence during infection, while also inducing a pro-inflammatory environment that enhances macrophage activation (Li *et al*., 2015; Leech *et al*., 2017). The increased pro-inflammatory state observed in macrophages in response to SSBP was associated with a significant reduction in the intramacrophagic survival of several virulent *S. aureus* strains, notably the highly pathogenic MRSA USA300. Moreover, our *in vivo* studies demonstrated that SSBP significantly reduced the bacterial burden, as evidenced by improved survival rates in zebrafish infection models. Altogether, these findings provide strong evidence that SSBP actively contributes to virulence attenuation *in vivo*, as both overexpression of the protein in *S. aureus* and its direct administration significantly enhanced zebrafish survival following infection.

### Roles and Mechanisms of SSBP

The mechanism of action of SSBP appears to involve modulation of the inflammatory response, highlighting its unique immunomodulatory capabilities that extend beyond its conventional role as a DNA-binding protein. Research on single-stranded DNA binding proteins has predominantly focused on Gram-negative bacteria, such as *E. coli*, while Gram-positive bacteria have received comparatively less attention (Antony and Lohman, 2019). SSBs are well-known for their essential roles in protecting ssDNA from nucleolytic degradation by binding with high affinity and without sequence specificity. However, recent studies have uncovered additional, often unexpected functions of SSBs, particularly in host-pathogen interactions. For instance, SSBX, an SSB produced by *Xanthomonas* bacteria, triggers hypersensitive responses in non-host plants, including oxidative bursts and the activation of MAPK and salicylic acid pathways (Li *et al*., 2013). Transferring the *ssb* gene from *Xanthomonas* to *Nicotiana benthamiana* enhanced plant growth and resistance to various stresses, including bacterial infections and salt stress (Cao *et al*., 2018). Similarly, targeting ssDNA with single-stranded DNA-binding proteins in *Neisseria gonorrhoeae* has been shown to disrupt biofilm formation, a critical virulence trait (Zweig *et al*., 2014).

Interestingly, while SSBP shares a high level of sequence conservation across prokaryotic and eukaryotic species, this does not preclude the possibility of unique functions. Within the ROSA-like prophage context, these functions may include specialized roles in modulating the immune response, particularly by enhancing macrophage activation and driving cytokine production. This is illustrated by the observed increase in IL-6 levels and the reduction in IL-10. While TNF-α was also elevated in infected macrophages, its expression did not significantly differ between conditions, suggesting that its regulation was not specifically influenced by SSBP. Such adaptations could reflect a unique evolutionary strategy that integrates immune modulation with conventional DNA-binding activities. Its role within the ROSA-like prophage context could involve modulating host immune responses, such as macrophage activation or cytokine production, alongside its potential DNA-binding activities. This dual functionality, combining conventional ssDNA protection with immune modulation, represents an adaptive trait that likely extends beyond its primary function.

It is important to note that while the generation of a clean *ssbP* knockout mutant would represent a more direct approach to assess its function, this strategy carries a high risk of polar effects due to the dense genomic organization of the prophage locus. The *ssbP* gene is closely flanked by other potentially co-transcribed genes, and its deletion could unintentionally disrupt their expression. To avoid this, we used the prophage-excised strain (ΔROSA) complemented with a plasmid expressing *ssbP*, which allowed us to isolate the contribution of this gene in a clean background. Although this strategy does not preserve the broader prophage context and may overlook potential interactions with other phage-encoded elements, the clear phenotypic changes observed upon *ssbP* complementation strongly support its independent role in modulating macrophage responses and bacterial clearance. Future approaches using refined genetic tools, such as CRISPR-based editing or transcriptional silencing, could allow specific targeting of *ssbP* while maintaining the integrity of the surrounding prophage, thus enabling a more precise dissection of its function within its native genomic context.

Moreover, dual RNA-Seq analyses conducted during macrophage infection revealed that while SSBP expression strongly influenced host immune gene expression, it did not significantly alter the transcriptional profile of bacterial virulence-associated genes. This indicates that the observed attenuation of bacterial survival is not due to altered bacterial gene expression but rather to host-mediated mechanisms, consistent with the secretion of SSBP and its role as an extracellular immunomodulatory factor. Notably, *S. aureus* has been shown to survive in macrophages outside the phagolysosomal compartment, and various staphylococcal proteins have been identified that reach the host cytosol to exert their effects (de Vor *et al*., 2020; Pidwill *et al*., 2021). SSBP might similarly act within these intracellular niches to alter host signaling pathways, although the precise mechanism of cellular entry or intracellular trafficking remains to be elucidated

One intriguing question arising from this study is why a colonizing strain of *S. aureus* would secrete a protein like SSBP that activates the immune system, potentially reducing its own survival and virulence. Several hypotheses could explain this paradox. First, activating a controlled immune response may help the bacteria establish a niche by suppressing overgrowth of competing microbes without triggering a strong enough response to eradicate *S. aureus*. Second, this strategy could reflect a balance between persistence and clearance, where a mild pro-inflammatory environment fosters tolerance rather than elimination, allowing the colonizing strain to coexist with the host. Third, this modulation might enable *S. aureus* to avoid overwhelming the host, which could lead to severe immune responses detrimental to both the bacterium and the host. Additionally, the ability to fine-tune the immune response may provide *S. aureus* with an advantage in adapting to different host conditions, including asymptomatic colonization and infection transitions. Moreover, *S. aureus* is known to form biofilms significantly faster, which further enhances its ability to evade phagocytosis and persist within the host. Finally, the ROSA-like prophage could confer selective advantages under specific conditions, such as chronic infections, where immune modulation benefits both the host and the colonizing strain by limiting invasive pathogens. Another possibility is that, under certain environmental stressors, excision of the prophage could allow the bacterium to shift from an asymptomatic carrier state to a more virulent infectious form. However, this scenario appears unlikely, as the ROSA-like prophage is highly stable and was particularly challenging to excise experimentally, suggesting that its presence is deeply integrated into the bacterial lifestyle. Moreover, the ROSA-like prophage lacks key lytic genes required for the production of functional phage particles, indicating that it is a defective prophage unable to enter a productive lytic cycle. Consistently, we never observed any lysis plaques or signs of virion production following macrophage infection in our conditions, supporting the view that excision and phage-mediated competition are not part of its functional repertoire. To date, NSA1385 is the only reported strain carrying this ROSA-like prophage with SSBP expression linked to reduced intracellular survival, and no similar phenotype has been described for SSB homologs in other *S. aureus* strains. Therefore, these evolutionary strategies highlight the complex interactions between *S. aureus*, its prophage-encoded proteins, and the host immune system, positioning this strain as a potential early example of bacterial adaptation to commensalism.

### Implications and Therapeutic Potential

In chronic infections, such as those that can occur in DFUs, the intracellular survival of *S. aureus* has been identified as a significant factor contributing to infection recurrence (Rasigade *et al*., 2016). Reduced susceptibility to antibiotics in such cases highlights the need for alternative strategies (Peyrusson *et al*., 2020). In this context, SSBP ability to modulate the immune response and enhance macrophage activation offers a novel and complementary approach to reducing intracellular bacterial survival and addressing persistent infections. Targeting intracellular survival mechanisms has shown promise. For example, silencing the human gene *tram2* with siRNA significantly reduced bacterial load without harming host cells (Bravo-Santano *et al*., 2019a). Similarly, tyrosine kinase inhibitors, such as Ibrutinib, Dasatinib, and Crizotinib, have demonstrated efficacy in reducing intracellular bacterial survival (Bravo-Santano *et al*., 2019b). Specific virulence factors, such as the acid phosphatase SapS (Ahmad-Mansour *et al*., 2022), tyrosine phosphatase PtpA (Gannoun-Zaki *et al*., 2018), and phosphoarginine phosphatase PtpB (Elhawy *et al*., 2021), have also been targeted to reduce intracellular persistence. Furthermore, bacteriophage-derived proteins, like those described by (Yang *et al*., 2018), effectively attenuate *S. aureus* virulence and enhance host immune defenses.

In the present study, SSBP ability to reduce the intracellular survival of *S. aureus* through immunomodulation provides a complementary approach. The ROSA-like prophage, previously shown to be stable within *S. aureus* (Messad *et al*., 2015), encodes SSBP, which plays a pivotal role in decreasing bacterial virulence. Overexpression of SSBP or addition of recombinant protein significantly reduced *S. aureus* survival in macrophages without inducing cytotoxicity, indicating a direct role in modulating host-pathogen interactions. SSBP potential to interfere with intracellular pathways and facilitate bacterial clearance positions it as a promising candidate for therapeutic development. Moreover, SSBP significantly reduced bacterial virulence, as evidenced by decreased pathogenicity and improved survival rates in the zebrafish infection model. This model provides valuable insights into host-pathogen interactions and the biological activity of SSBP, but further studies in mammalian models, such as mice, will be essential to confirm these results and assess their translational relevance to human infections. However, while these findings highlight the therapeutic potential of SSBP, it is crucial to consider the potential activation of the “double-edged sword” mechanism (Ashida *et al*., 2011). In the case of the NSA1385 strain harboring the phage and thus SSBP, the strain adopts a colonizing strategy and does not trigger this detrimental mechanism, likely due to the fine-tuned regulation of SSBP expression. Nevertheless, this aspect warrants further investigation, particularly if recombinant pro-inflammatory SSBP is to be considered as a therapeutic agent, to ensure it does not inadvertently exacerbate inflammation or tissue damage.

## CONCLUSION

The ability of *S. aureus* to survive within host cells is a critical factor in chronic infections, including those associated with DFUs. The dual effects of SSBP on *S. aureus* pathogenicity—attenuating virulence while promoting immune clearance—underscore its potential as a novel anti-virulence agent. This study highlights SSBP’s unique capacity to stimulate a pro-inflammatory immune response, which facilitates bacterial clearance while minimizing host cytotoxicity. These qualities position SSBP as a promising candidate for combating antibiotic-resistant *S. aureus* and other Gram-positive pathogens. Future investigations should focus on its therapeutic applications, such as its integration into treatment strategies for intracellular infections or its use as an immunomodulatory agent to enhance existing therapies. Nonetheless, further research is essential to clarify its precise mechanisms of action and fully assess its therapeutic potential.

## Author Contributions

Conceptualization, J.P.L. and V.M.; methodology, N.A.M., C.N.A.N., M.M., L.P., S.H-B., V.M., and J.P.L.; software, M.Mo., N.A.M., and M.M.; validation, V.M., and J.P.L.; formal analysis, N.A.M., C.N.A.N., M.M., L.P., S.H-B.; investigation, N.A.M., C.N.A.N., M.M., L.P., S.H-B.; resources, V.M. and J.P.L.; writing—original draft preparation, N.A.M., V.M., and J.P.L.; writing—review and editing, N.A.M., C.N.A.N., M.M., L.P., S.H-B, M.Mo., C.D.R., A.S., V.M., and J.P.L.; supervision, V.M. and J.P.L.; project administration, V.M. and J.P.L.; funding acquisition, V.M. and J.P.L.

## Financial Support and Acknowledgments

N.A.M. and C.N.A.N. received PhD grants from InfectioPôle Sud Méditerranée. C.D.R., A.S., and J.P.L. were supported by the University Hospital of Nîmes, which provided structural, human, and financial support through the “Thématiques phares” program.

## REFERENCES

Ahmad-Mansour, N., Elhawy, M.I., Huc-Brandt, S., Youssouf, N., Pätzold, L., Martin, M., et al. (2022) Characterization of the Secreted Acid Phosphatase SapS Reveals a Novel Virulence Factor of *Staphylococcus aureus* That Contributes to Survival and Virulence in Mice. Int J Mol Sci 23: 14031.

Ahmad-Mansour, N., Plumet, L., Pouget, C., Kissa, K., Dunyach-Remy, C., Sotto, A., et al. (2023) The ROSA-Like Prophage Colonizing *Staphylococcus aureus* Promotes Intracellular Survival, Biofilm Formation, and Virulence in a Chronic Wound Environment. J Infect Dis jiad218.

Antony, E., Weiland, E., Yuan, Q., Manhart, C.M., Nguyen, B., Kozlov, A.G., et al. (2013) Multiple C-Terminal Tails within a Single *E. coli* SSB Homotetramer Coordinate DNA Replication and Repair. J Mol Biol 425: 4802–4819.

Ashida, H., Mimuro, H., Ogawa, M., Kobayashi, T., Sanada, T., Kim, M., and Sasakawa, C. (2011) Cell death and infection: a double-edged sword for host and pathogen survival. J Cell Biol 195: 931–942.

Baba, T., Bae, T., Schneewind, O., Takeuchi, F., and Hiramatsu, K. (2008) Genome Sequence of *Staphylococcus aureus* Strain Newman and Comparative Analysis of Staphylococcal Genomes: Polymorphism and Evolution of Two Major Pathogenicity Islands. J Bacteriol 190: 300–310.

Bae, T., Baba, T., Hiramatsu, K., and Schneewind, O. (2006) Prophages of *Staphylococcus aureus* Newman and their contribution to virulence. Mol Microbiol 62: 1035–1047.

Boukamp, P., Petrussevska, R.T., Breitkreutz, D., Hornung, J., Markham, A., and Fusenig, N.E. (1988) Normal keratinization in a spontaneously immortalized aneuploid human keratinocyte cell line. J Cell Biol 106: 761–771.

Bravo-Santano, N., Capilla-Lasheras, P., Mateos, L.M., Calle, Y., Behrends, V., and Letek, M. (2019a) Identification of novel targets for host-directed therapeutics against intracellular *Staphylococcus aureus*. Sci Rep 9: 15435.

Bravo-Santano, N., Stölting, H., Cooper, F., Bileckaja, N., Majstorovic, A., Ihle, N., et al. (2019b) Host-directed kinase inhibitors act as novel therapies against intracellular *Staphylococcus aureus*. Sci Rep 9: 4876.

Cao, Y., Yang, M., Ma, W., Sun, Y., and Chen, G. (2018) Overexpression of SSBXoc, a Single-Stranded DNA-Binding Protein From *Xanthomonas oryzae* pv. *oryzicola*, Enhances Plant Growth and Disease and Salt Stress Tolerance in Transgenic *Nicotiana benthamiana*. Front Plant Sci 9: 953.

Charbonnier, Y., Gettler, B., François, P., Bento, M., Renzoni, A., Vaudaux, P., et al. (2005) A generic approach for the design of whole-genome oligoarrays, validated for genomotyping, deletion mapping and gene expression analysis on *Staphylococcus aureus*. BMC Genomics 6: 95.

Cheung, G.Y.C., Bae, J.S., and Otto, M. (2021) Pathogenicity and virulence of *Staphylococcus aureus*. Virulence 12: 547–569.

Corona, R.I., and Guo, J. (2016) Statistical analysis of structural determinants for protein-DNA binding specificity. Proteins 84: 1147–1161.

Diep, B.A., Gill, S.R., Chang, R.F., Phan, T.H., Chen, J.H., Davidson, M.G., et al. (2006) Complete genome sequence of USA300, an epidemic clone of community-acquired meticillin-resistant *Staphylococcus aureus*. Lancet Lond Engl 367: 731–739.

Elhawy, M.I., Huc-Brandt, S., Pätzold, L., Gannoun-Zaki, L., Abdrabou, A.M.M., Bischoff, M., and Molle, V. (2021) The Phosphoarginine Phosphatase PtpB from *Staphylococcus aureus* Is Involved in Bacterial Stress Adaptation during Infection. Cells 10: 645.

Feiner, R., Argov, T., Rabinovich, L., Sigal, N., Borovok, I., and Herskovits, A.A. (2015) A new perspective on lysogeny: prophages as active regulatory switches of bacteria. Nat Rev Microbiol 13: 641–650.

Fournier, B., and Philpott, D.J. (2005) Recognition of *Staphylococcus aureus* by the innate immune system. Clin Microbiol Rev 18: 521–540.

Fraunholz, M., and Sinha, B. (2012) Intracellular *Staphylococcus aureus*: live-in and let die. Front Cell Infect Microbiol 2: 43.

Gannoun-Zaki, L., Pätzold, L., Huc-Brandt, S., Baronian, G., Elhawy, M.I., Gaupp, R., et al. (2018) PtpA, a secreted tyrosine phosphatase from *Staphylococcus aureus*, contributes to virulence and interacts with coronin-1A during infection. J Biol Chem 293: 15569–15580.

Ge, S.X., Jung, D., and Yao, R. (2020) ShinyGO: a graphical gene-set enrichment tool for animals and plants. Bioinformatics 36: 2628–2629.

Gunaratnam, G., Tuchscherr, L., Elhawy, M.I., Bertram, R., Eisenbeis, J., Spengler, C., et al. (2019) ClpC affects the intracellular survival capacity of *Staphylococcus aureus* in non-professional phagocytic cells. Sci Rep 9: 16267.

Guo, J.-T., and Malik, F. (2022) Single-Stranded DNA Binding Proteins and Their Identification Using Machine Learning-Based Approaches. Biomolecules 12: 1187.

Hiramatsu, K., Ito, T., Tsubakishita, S., Sasaki, T., Takeuchi, F., Morimoto, Y., et al. (2013) Genomic Basis for Methicillin Resistance in *Staphylococcus aureus*. Infect Chemother 45: 117–136.

Holden, M.T.G., Feil, E.J., Lindsay, J.A., Peacock, S.J., Day, N.P.J., Enright, M.C., et al. (2004) Complete genomes of two clinical *Staphylococcus aureus* strains: evidence for the rapid evolution of virulence and drug resistance. Proc Natl Acad Sci U S A 101: 9786–9791.

Hommes, J.W., and Surewaard, B.G.J. (2022) Intracellular Habitation of *Staphylococcus aureus*: Molecular Mechanisms and Prospects for Antimicrobial Therapy. Biomedicines 10: 1804.

Lebeurre, J., Dahyot, S., Diene, S., Paulay, A., Aubourg, M., Argemi, X., et al. (2019) Comparative Genome Analysis of *Staphylococcus lugdunensis* Shows Clonal Complex-Dependent Diversity of the Putative Virulence Factor, ess/Type VII Locus. Front Microbiol 10: 2479.

Leech, J.M., Lacey, K.A., Mulcahy, M.E., Medina, E., and McLoughlin, R.M. (2017) IL-10 Plays Opposing Roles during *Staphylococcus aureus* Systemic and Localized Infections. J Immunol Author Choice 198: 2352–2365.

Li, Y.-R., Ma, W.-X., Che, Y.-Z., Zou, L.-F., Zakria, M., Zou, H.-S., and Chen, G.-Y. (2013) A Highly-Conserved Single-Stranded DNA-Binding Protein in *Xanthomonas* Functions as a Harpin-Like Protein to Trigger Plant Immunity. PLOS ONE 8: e56240.

Li, Z., Peres, A.G., Damian, A.C., and Madrenas, J. (2015) Immunomodulation and Disease Tolerance to *Staphylococcus aureus*. Pathogens 4: 793–815.

Löffler, B., Tuchscherr, L., Niemann, S., and Peters, G. (2014) *Staphylococcus aureus* persistence in non-professional phagocytes. Int J Med Microbiol 304: 170–176.

Malachowa, N., and DeLeo, F.R. (2010) Mobile genetic elements of *Staphylococcus aureus*. Cell Mol Life Sci CMLS 67: 3057–3071.

Marceau, A.H. (2012) Functions of single-strand DNA-binding proteins in DNA replication, recombination, and repair. Methods Mol Biol Clifton NJ 922: 1–21.

Messad, N., Prajsnar, T.K., Lina, G., O’Callaghan, D., Foster, S.J., Renshaw, S.A., et al. (2015) Existence of a Colonizing *Staphylococcus aureus* Strain Isolated in Diabetic Foot Ulcers. Diabetes 64: 2991–2995.

Monk, I.R., Tree, J.J., Howden, B.P., Stinear, T.P., and Foster, T.J. (2015) Complete Bypass of Restriction Systems for Major *Staphylococcus aureus* Lineages. mBio 6 https://www.ncbi.nlm.nih.gov/pmc/articles/PMC4447248/. Accessed May 31, 2023.

Peyrusson, F., Varet, H., Nguyen, T.K., Legendre, R., Sismeiro, O., Coppée, J.-Y., et al. (2020) Intracellular *Staphylococcus aureus* persisters upon antibiotic exposure. Nat Commun 11: 2200.

Pidwill, G.R., Gibson, J.F., Cole, J., Renshaw, S.A., and Foster, S.J. (2021) The Role of Macrophages in *Staphylococcus aureus* Infection. Front Immunol 11 https://www.frontiersin.org/article/10.3389/fimmu.2020.620339. Accessed June 21, 2022.

Rasigade, J.-P., Dunyach-Rémy, C., Sapin, A., Messad, N., Trouillet-Assant, S., Dupieux, C., et al. (2016) A Prophage in Diabetic Foot Ulcer-Colonizing *Staphylococcus aureus* Impairs Invasiveness by Limiting Intracellular Growth. J Infect Dis 214: 1605–1608.

Scherl, A., François, P., Charbonnier, Y., Deshusses, J.M., Koessler, T., Huyghe, A., et al. (2006) Exploring glycopeptide-resistance in *Staphylococcus aureus*: a combined proteomics and transcriptomics approach for the identification of resistance-related markers. BMC Genomics 7: 296.

Schwendener, S., and Perreten, V. (2015) New Shuttle Vector-Based Expression System To Generate Polyhistidine-Tagged Fusion Proteins in *Staphylococcus aureus* and *Escherichia coli*. Appl Environ Microbiol 81: 3243–3254.

Secor, P.R., James, G.A., Fleckman, P., Olerud, J.E., McInnerney, K., and Stewart, P.S. (2011) *Staphylococcus aureus* Biofilm and Planktonic cultures differentially impact gene expression, mapk phosphorylation, and cytokine production in human keratinocytes. BMC Microbiol 11: 143.

Sudo, H., Kodama, H.A., Amagai, Y., Yamamoto, S., and Kasai, S. (1983) *In vitro* differentiation and calcification in a new clonal osteogenic cell line derived from newborn mouse calvaria. J Cell Biol 96: 191–198.

Takashiba, S., Van Dyke, T.E., Amar, S., Murayama, Y., Soskolne, A.W., and Shapira, L. (1999) Differentiation of Monocytes to Macrophages Primes Cells for Lipopolysaccharide Stimulation via Accumulation of Cytoplasmic Nuclear Factor κB. Infect Immun 67: 5573–5578.

Tong, S.Y.C., Davis, J.S., Eichenberger, E., Holland, T.L., and Fowler, V.G. (2015) *Staphylococcus aureus* Infections: Epidemiology, Pathophysiology, Clinical Manifestations, and Management. Clin Microbiol Rev 28: 603–661.

Tsuchiya, S., Yamabe, M., Yamaguchi, Y., Kobayashi, Y., Konno, T., and Tada, K. (1980) Establishment and characterization of a human acute monocytic leukemia cell line (THP-1). Int J Cancer 26: 171–176.

Tuffs, S.W., Haeryfar, S.M.M., and McCormick, J.K. (2018) Manipulation of Innate and Adaptive Immunity by Staphylococcal Superantigens. Pathog Basel Switz 7: 53.

Vestergaard, M., Frees, D., and Ingmer, H. (2019) Antibiotic Resistance and the MRSA Problem. Microbiol Spectr 7: 7.2.18.

Vor, L. de, Rooijakkers, S.H.M., and Strijp, J.A.G. van (2020) Staphylococci evade the innate immune response by disarming neutrophils and forming biofilms. FEBS Lett 594: 2556–2569.

Yang, H., Xu, J., Li, W., Wang, S., Li, J., Yu, J., et al. (2018) *Staphylococcus aureus* virulence attenuation and immune clearance mediated by a phage lysin-derived protein. EMBO J 37: e98045.

Zweig, M., Schork, S., Koerdt, A., Siewering, K., Sternberg, C., Thormann, K., et al. (2014) Secreted single-stranded DNA is involved in the initial phase of biofilm formation by Neisseria gonorrhoeae. Environ Microbiol 16: 1040–1052.

