## Supplementary figures for "The *Staphylococcus aureus* prophage-encoded SSBP attenuates virulence and enhances IL-6-mediated macrophage clearance"

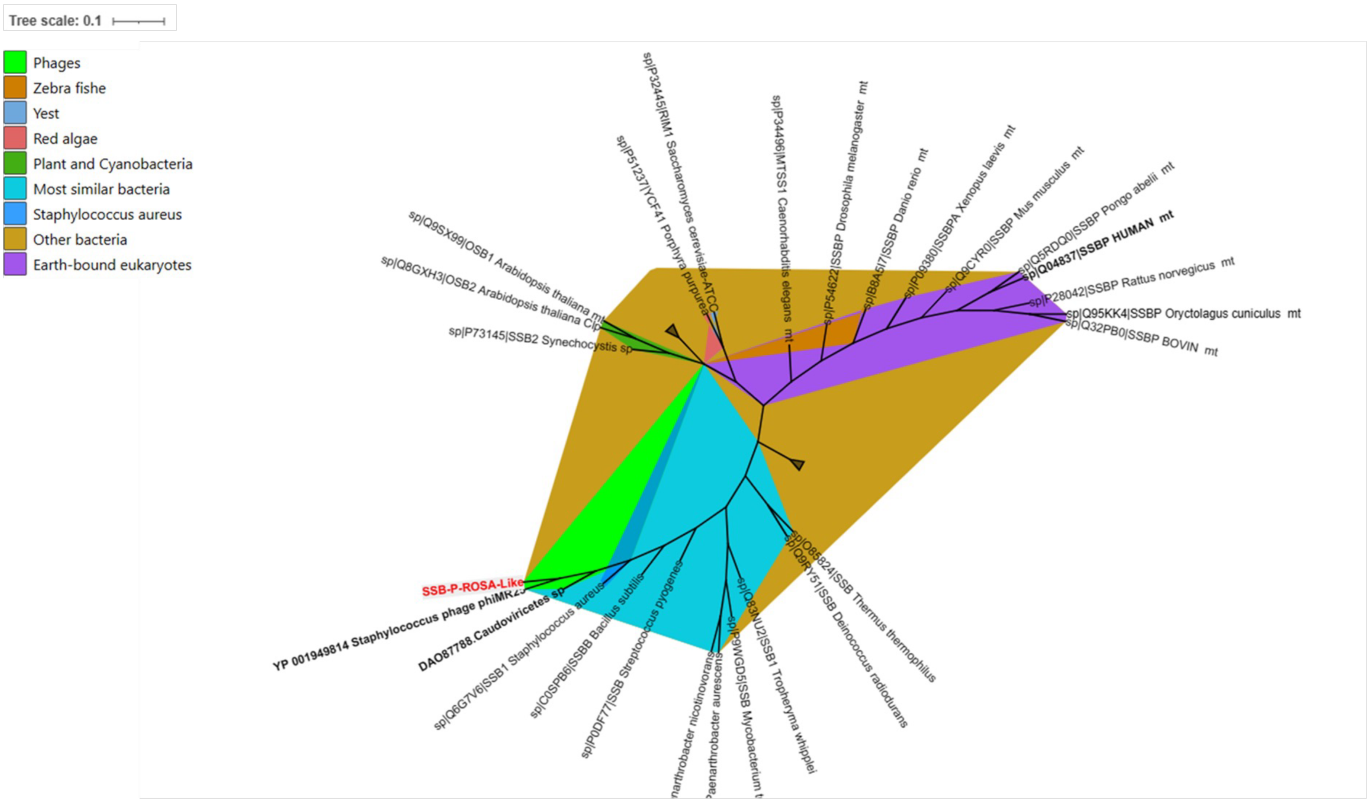


**Supplementary Figure 1. Phylogenetic analysis of ROSA-like single-stranded DNA-binding protein (SSBP).** The phylogenetic tree was constructed using the ROSA-like single-stranded DNA-binding protein (SSBP) from *Staphylococcus aureus* and 51 homologous proteins retrieved from the UniProt database. These homologues include proteins from phages, bacteria, cyanobacteria, mitochondrial proteins from protozoa, plants, and animals (including the human mitochondrial single-stranded DNA-binding protein), as well as the *Arabidopsis thaliana* chloroplastic OSB2 protein and proteins from marine algae.

Sequence alignment was performed using the MAFFT online tool (available at MAFFT), and the phylogenetic tree was visualized using iTOL (Interactive Tree of Life). Notably, the ROSA-like SSBP protein exhibits close similarity to the *S. aureus* phage phiMR25 protein (GenBank accession no. YP_001949814), classified under *Caudoviricetes* sp., and shares high similarity with the *S. aureus* SSB1 protein, which is known for its role in single-stranded DNA binding.

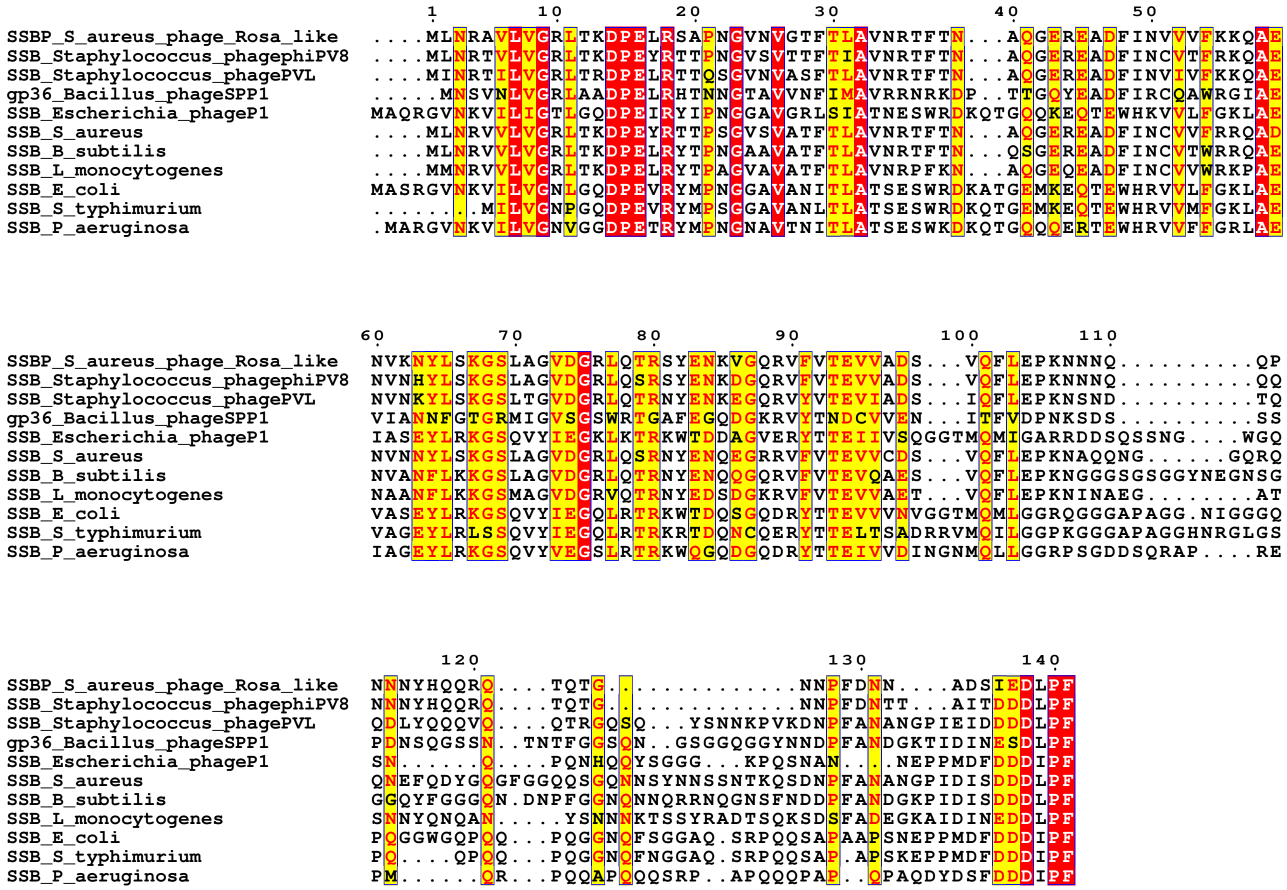


**Supplementary Figure 2. Multiple sequence alignment of single-stranded DNA-binding proteins from bacterial and bacteriophage origins**. The alignment includes sequence identifiers comprising the protein name and source organism. Conserved residues are highlighted in red, indicating their potential functional or structural importance. The alignment was generated using the ESPript server (Robert, X., and Gouet, P. 2014. Deciphering key features in protein structures with the new ENDscript server. Nucleic Acids Res 42: W320-324), providing a clear visualization of conserved motifs.

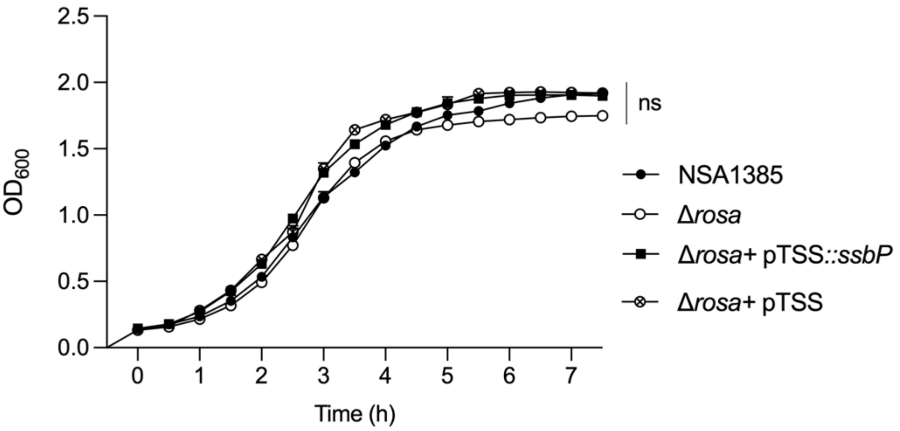


**Supplementary Figure 3. Growth kinetics of *S. aureus* NSA1385, Δ*rosa*, Δ*rosa* + pTSS*::ssbP* and Δ*rosa* + pTSS (negative control) in tryptic soy broth (TSB).** Cultures were grown in 96-well plates at 37°C with shaking at 108 rpm, using a microplate reader (Tecan, Model Spark, Grödig, Austria GmbH) . Data are presented as the mean optical density at 600 nm ($OD_{600}$) readings ± standard deviation (n = 3). At certain time points, error bars are too narrow to be visible.
