## Supplementary tables for "The *Staphylococcus aureus* prophage-encoded SSBP attenuates virulence and enhances IL-6-mediated macrophage clearance"

**Supplementary Table 1.** Strains and plasmids used in this study

| Strain | Description | References |
|---|---|---|
| ***S. aureus*** | | |
| NSA1385 | Clinical strain isolated from colonized Diabetic Foot Ulcer (DFU) (grade 1)* | (1) |
| Δ*rosa* | The ΔROSA corresponds to a prophage-free derivative of NSA1385 obtained after excision of the ROSA-like prophage | (1) |
| Δ*rosa*+ pTSS::*ssbP* | Δ*rosa* complemented with pTSS:: *ssbP* plasmid | This study |
| NSA739 | Clinical strain isolated from colonized Diabetic foot ulcer (DFU) (grade 3) | (1) |
| SA564 | Clinical isolate | (2) |
| USA300 | Methicillin-resistant *S. aureus* strain | (3) |
| ***E. coli*** | | |
| BL21 star (DE3) | *E. coli* strain allowing a high-level recombinant protein expression. IPTG-inducible T7 RNA polymerase | Invitrogen |
| **Plasmid** | | |
| pTSSCm-$P_{cap}$ | *E. coli- S. aureus* shuttle vector for native expression | (4) |
| pTSS*::ssb-P* | pTSS derivative used to express C-terminal Spot- tagged fusion of ROSA-like SSBP | This study |
| pET19b | *E. coli* vector for IPTG inducible protein expression*; bla* | Novagen |
| pET19b::*ssbP* | pET19b derivative used to express His-tagged fusion of ROSA-like SSBP; *bla* | This study |

*IWGDF classification of foot infection (https://iwgdfguidelines.org/wp-content/uploads/2023/07/IWGDF-2023-03-Classification-Guideline.pdf)

**Supplementary Table 2.** Primers used in this study

| Primer | Direction | Sequence (5′-3′) |
|---|---|---|
| #103 | For. | GAGGATCTTCCTTTTGGGTCATCAGGGCCAGATCG |
| #54 | Rev. | TGATACTGCACGAACACGATC |
| #99 | For. | GTTGGCCGATTCATTAATGCAG |
| #104 | Rev. | TCGAGCATGCGGATCCTAAGAACTCCAATGTGATA |
| #101 | For. | TAGCTGCAGGAATTCATGTTAAACAGAGCAGTATT |
| #102 | Rev. | TGGCCCTGATGACCCAAAAGGAAGATCCTCTATAG |
| #129 | For. | TTCCAGGGCCATATGTTAAACAGAGCAGTATTAGT |
| #130 | Rev. | TAGTTATTAGGATCCCTAAAAAGGAAGATCCTCTA |


1. Messad N, Prajsnar TK, Lina G, O'Callaghan D, Foster SJ, Renshaw SA, Skaar EP, Bes M, Dunyach-Remy C, Vandenesch F, Sotto A, Lavigne J-P. 2015. Existence of a Colonizing *Staphylococcus aureus* Strain Isolated in Diabetic Foot Ulcers. Diabetes 64:2991–2995.

2. Somerville GA, Beres SB, Fitzgerald JR, DeLeo FR, Cole RL, Hoff JS, Musser JM. 2002. *In vitro* serial passage of *Staphylococcus aureus*: changes in physiology, virulence factor production, and agr nucleotide sequence. J Bacteriol 184:1430–1437.

3. Tenover FC, Goering RV. 2009. Methicillin-resistant *Staphylococcus aureus* strain USA300: origin and epidemiology. Journal of Antimicrobial Chemotherapy 64:441–446.

4. Schwendener S, Perreten V. 2015. New Shuttle Vector-Based Expression System To Generate Polyhistidine-Tagged Fusion Proteins in *Staphylococcus aureus* and *Escherichia coli*. Applied and Environmental Microbiology 81:3243–3254.
